# In-vitro Metabolite Identification for MEDI7219, an Oral GLP-1 Peptide, using LC-MS/MS with CID and EAD Fragmentation

**DOI:** 10.1101/2024.07.26.605352

**Authors:** Kate Liu, Yue Huang, Taoqing Wang, Ruipeng Mu, Anton I. Rosenbaum

## Abstract

Oral peptide therapeutics typically suffer from short half-lives as they are readily degraded by digestive enzymes. Systematic peptide engineering along with formulation optimization led to the development of a clinical candidate MEDI7219, an orally-bioavailable GLP-1 peptide, that is much more stable than wild type GLP-1 or semaglutide. In this study, we elucidated peptide biotransformation products using in vitro pancreatin assay that employed both collision-induced dissociation (CID) and electron-activated dissociation (EAD) LC-MS/MS methods. Using this approach, we have confidently identified a total of 13 metabolites. Relative quantification of these metabolites over time showed sequential cleavage pattern as peptides were further digested to smaller fragments. These 13 metabolites mapped to 8 cleavage sites on MEDI7219 structure. Most of these cleavage sites can be explained by the specificity of digestive enzymes, e.g. ,trypsin, pepsin and elastase. α-methyl-L-phenylalanine appeared to be well protected from chymotrypsin and pepsin digestion since no cleavage peptides ending with α-methyl-L-phenylalanine were observed. These study results expand upon previously published stability data and provide new insights on potential GLP1 proteolytic liabilities for future engineering. Furthermore, this study exemplifies the application of pancreatin in vitro system methodology as a valuable tool for understanding metabolism of oral peptide therapeutics *in vitro*. Additionally, orthogonal MS fragmentation modes offered improved confidence in identification for peptide unknown metabolites.

**Significance Statement:** In vitro metabolite identification of oral GLP1 peptide that uncovers potential proteolytic hotspots that can inform future oral peptide engineering.

## Introduction

Peptide therapeutics have been increasingly used in treatment of metabolic and chronic diseases such as diabetes (Fosgerau & Hoffmann, 2015) and GLP-1 receptor agonists are established therapeutics for patients with type 2 diabetes and obesity. However, administration by subcutaneous injection can reduce patient compliance. Oral delivery of peptide therapeutics offers greater convenience for long term treatment, but it is challenged by low gastrointestinal (GI) permeability and high susceptibility to degradation by GI and serum proteases. Oral peptides tend to suffer from poor bioavailability and short circulating half-lives. To overcome these challenges, glucagon-like peptide 1 (GLP-1) receptor agonist was specifically engineered for digestion resistance and formulated for oral delivery resulting in MEDI7219 (Pechenov et al., 2021; Tyagi et al., 2023; Tyagi et al., 2021).

GLP-1 is a peptide hormone that stimulates glucose-dependent insulin release leading to improved glycemic control and delays gastric emptying. Wild type GLP-1 is highly susceptible to enzymatic degradation with plasma half-life of a few minutes. Circulating GLP-1 is rapidly degraded by serum proteases including dipeptidyl peptidase-4 (DPP-IV) and neprilysin and digestive proteases such as pepsin, trypsin, chymotrypsin and elastase. These enzymes digest peptides into di-and tri-peptides presenting a challenge in GI stability.

Semaglutide was the first approved oral GLP-1 receptor agonist (Mahapatra et al., 2022) and it incorporated peptide engineering approaches such as unnatural amino acid substitution to protect proteolytic sites, and lipidation to promote plasma protein binding to reduce renal clearance. However, semaglutide is still rapidly degraded in GI tract (Pechenov et al., 2021).

Systematic multi-disciplinary approach to engineering led to rational selection of the clinical lead, MEDI7219, a GLP-1 peptide with bis-lipidation and 12 amino acid substitutions and 5 α- methyl modifications (Pechenov et al., 2021). This peptide was potent as well as proteolytically more stable than the wild type or semaglutide. In vitro study in FaSSIF/pancreatin showed over 60 % of MEDI7219 remained intact after 2 h (Pechenov et al., 2021). This measurement was done using HPLC with MEDI7219 detection by UV absorption at 210 nm. In this study, we identified key metabolites from MEDI7219 in pancreatin in vitro assay using both collision-induced dissociation (CID) and electron-activated dissociation (EAD) fragmentation methods with LC-MS/MS. These metabolites reveal vulnerabilities in peptide structure that can inform future peptide engineering. As the field of therapeutic peptides grows, characterizing peptide biotransformation will be increasingly valuable to understand safety and stability of drug candidates. Unlike small molecule drug development where metabolite identification (metID) is more routine, peptide metabolism remains less explored, highlighting the need to build knowledge in this area (Cuyckens et al., 2012; Yu et al., 2020).

## Materials and Methods

### Materials and Reagents

MEDI7219 was provided by AstraZeneca (Gaithersburg, MD). Pancreatin from porcine pancreas (8X USP grade, P7545) was purchased from Sigma Aldrich (St. Louis, MO). Phosphate-buffered saline (PBS) was purchased from Thermo Fisher Scientific (Waltham, MA). Acetonitrile was from Millipore Sigma (Burlington, MA). 0.1 % formic acid was purchased from Thermo Fisher Scientific (Waltham, MA).

### Structural Characterization of MEDI7219 peptide by CID and EAD

MEDI7219 solution (10 µg/mL) in PBS was injected on SCIEX Exion LC system coupled with SCIEX ZenoTOF 7600 mass spectrometer. A 40 µL sample was loaded onto Waters ACQUITY UPLC BEH C18 Column (PN186002350) for each injection at flow rate of 0.5 ml/min using mobile phases A (0.1 % formic acid in water) and B (0.1 % formic acid in acetonitrile). The separation was carried out with LC gradient of 35% B to 80% B within 4 min.

Data were acquired in positive DDA (data dependent acquisition) mode using either CID or EAD fragmentation. For CID method, the collision energy was 40 V and declustering potential was 80V. For EAD method, key parameters decided after optimization are electron KE (kinetic energy) 11 eV, current 5 µA and reaction time 20 ms. Data analysis was performed with SCIEX PeakView (v 2.2) with Biotools function by entering precursor peptide sequence and matching fragment ions (b/y/c/z as well as internal fragment ions y|a and y|b).

### In vitro incubation with pancreatin

MEDI7219 solution (10 µg/ml) was incubated with pancreatin (3 mg/ml) in pre-warmed PBS buffer at 37 °C for 0.5 h, 1 h, 2 h, 4 h and 6 h. A negative control sample was prepared as pancreatin in PBS buffer without MEDI7219 spiked in. A time zero sample was prepared as MEDI7219 in PBS without pancreatin. At each time point, 100 µl was sampled and quenched with 200 µl of acetonitrile. The quenched solution was centrifuged at 6000 *g* for 10 min. Supernatant was diluted 5-fold in mobile phase A (0.1 % formic acid in water) and 40 µl was injected to LC-MS/MS. Same LC-MS system was used here as structural characterization experiment. A longer LC gradient was applied where 20 % B was increased to 60 % B within 8 min.

For MS method, DDA (data dependent acquisition) experiment was used where ions in range of 800-1250 Da were selected for MS/MS. Same key settings for CID and EAD established in the structural characterization experiment described above were applied in MS method here.

Data were analyzed using Molecule Profiler beta version 1.3.2 (MP, SCIEX) beta version. Raw data from incubation at each time point was analyzed against the pancreatin blank sample in the MP software. Only peaks that were absent or significantly smaller in the blank sample were considered for identification. Amino acid sequence for MEDI7219 was entered as a processing parameter. Non-natural amino acids including α-methyl-L-alanine, α-methyl-L-phenylalanine, α-methyl-L-serine, α-methyl-L-lysine, as well as side-chain lipid on lysine (C35H63N5O11) were added as custom elements by providing molecular formula in the software. Peptide method was used to match all peptide fragment peaks with full lipid side chain. Both MS1 and MS2 were used for peak finding based on mass tolerance of 15 ppm. For MS parameters, charge states 2-5 were included in the search. For MS/MS parameters, theoretical b, y, c’, z· as well as y|a and y|b internal fragment ions were searched with mass tolerance of 15 ppm. Default Biologics biotransformation set was used in the processing method. For quantification, all isotope peaks as part of a cluster and multiple charge states of the same peptide were summed for peak area. Results from individual timepoints were consolidated in Correlation analysis with 0.1 min retention time (RT) tolerance and 15 ppm *m/z* tolerance to yield the final list of metabolites and peak areas from all samples.

## Results

### Structural Characterization of MEDI7219 peptide using CID vs EAD

CID is the most widely adopted MS/MS fragmentation method. EAD represents a recent advancement in electron-based fragmentation techniques that encompassing various mechanisms such as electron capture dissociation (ECD), hot ECD and electron-impact excitation of ions from organics (EIEIO) (Baba et al., 2021). Unlike traditional ECD or EIEIO, EAD device features a tunable kinetic energy electron beam allowing analysis of both small (singly charged) and large (multiply-charged) molecules on a single instrument, SCIEX ZenoTOF 7600 system (Baba et al., 2021). EAD can offer unique advantages compared to CID in structural elucidation of lipids (Calabrese et al., 2024), differentiation of isomers (Yang et al., 2024), and identification of labile post-translational modifications (Bons et al., 2023).

For EAD method, critical parameters evaluated using MEDI7219 included kinetic energy (KE) that ranged from 5 to 11 eV, current that ranged from 3 µA to 7 µA, and reaction time between 20 ms and 80 ms. We found that the higher the electron KE, the greater the intensities for EAD fragment c/z peaks (Figure S1, Figure S2). Current did not substantially impact EAD fragmentation (Figure S3, Figure S4). Reaction time could affect absolute intensities of fragments. The shorter the EAD reaction time, the higher the intensities of both fragments and parent (Figure S5, Figure S6). After this evaluation, final method employed the electron KE 11 eV, current 5 µA and reaction time 20 ms.

Rich fragmentation information of MEDI7219 was obtained in both CID and EAD mode (**Error! Reference source not found.**). CID resulted in better fragmentation peaks assignment than EAD, but both showed good sequence coverage. For CID, consecutive b/y ions were observed. For EAD, c/z ions dominate the spectrum. Both CID and EAD resulted in lipid side chain fragmentation, however the fragmentation positions were different. The prominent lipid fragments in CID were m/z 355 and m/z 500, whereas prominent lipid fragments in EAD were m/z 417 and m/z 562. In CID, carbon-oxygen bond at β position to N-H was more susceptible to fragmentation. Sine EAD fragmentation is guided by generating more stable radicals, breaking C-O bond next to amide carbonyl group would give a stabilized radical. These signature fragment ions can potentially be used to confirm identity of metabolites that contain lipid side chain, and CID and EAD complement each other enhancing identification confidence.

### Metabolite identification from enzymatic incubation

Pancreatin was chosen for the in vitro evaluation because the site of maximal gastrointestinal absorption for MEDI7219 oral peptide was small intestine and proximal colon (Pechenov et al., 2021). The original experiment performed in Pechenov et al. used pancreatin assay in fasted-state simulated intestinal fluid (FaSSIF). Pancreatin is the main proteolytic enzyme of the duodenum and it is commonly applied as in vitro model to mimic digestion in vivo (Seo et al., 2022). FaSSIF simulate the pH during fasted state in duodenum and it also contains bile salts and phospholipids that are important for evaluating dissolution of a drug or drug solubilization (Galia et al., 1998). For this study, we used PBS (pH 7.4) to simulate the small intestine environment (pH 6-7.4) where pancreatin is active, as solubility was expected to be comparable in FaSSIF and in a simpler buffer system like PBS.

After incubation of MEDI7219 with pancreatin in PBS for up to 6 hours, we identified a total of 13 metabolites (**Error! Reference source not found.**). Both CID and EAD fragmentation identified the same 13 metabolites confirming the results using two orthogonal fragmentation approaches. Confident assignment of these metabolites is based on precursor mass accuracy, precursor isotope distribution and MS/MS fragmentation, whenever available.

An example of metabolite identification is shown in Figure 2 and full identification assignments for all metabolites are provided in supplementary information (**Error! Reference source not found.**, Figure S7-Figure S19). The charge state was assigned based on the isotope cluster in MS1 (Figure 2A), and the structure of metabolite was confirmed based on the exact masses of precursor (Figure 2A) and product ions from CID and EAD (Figure 2B, C).

**Figure 1.**
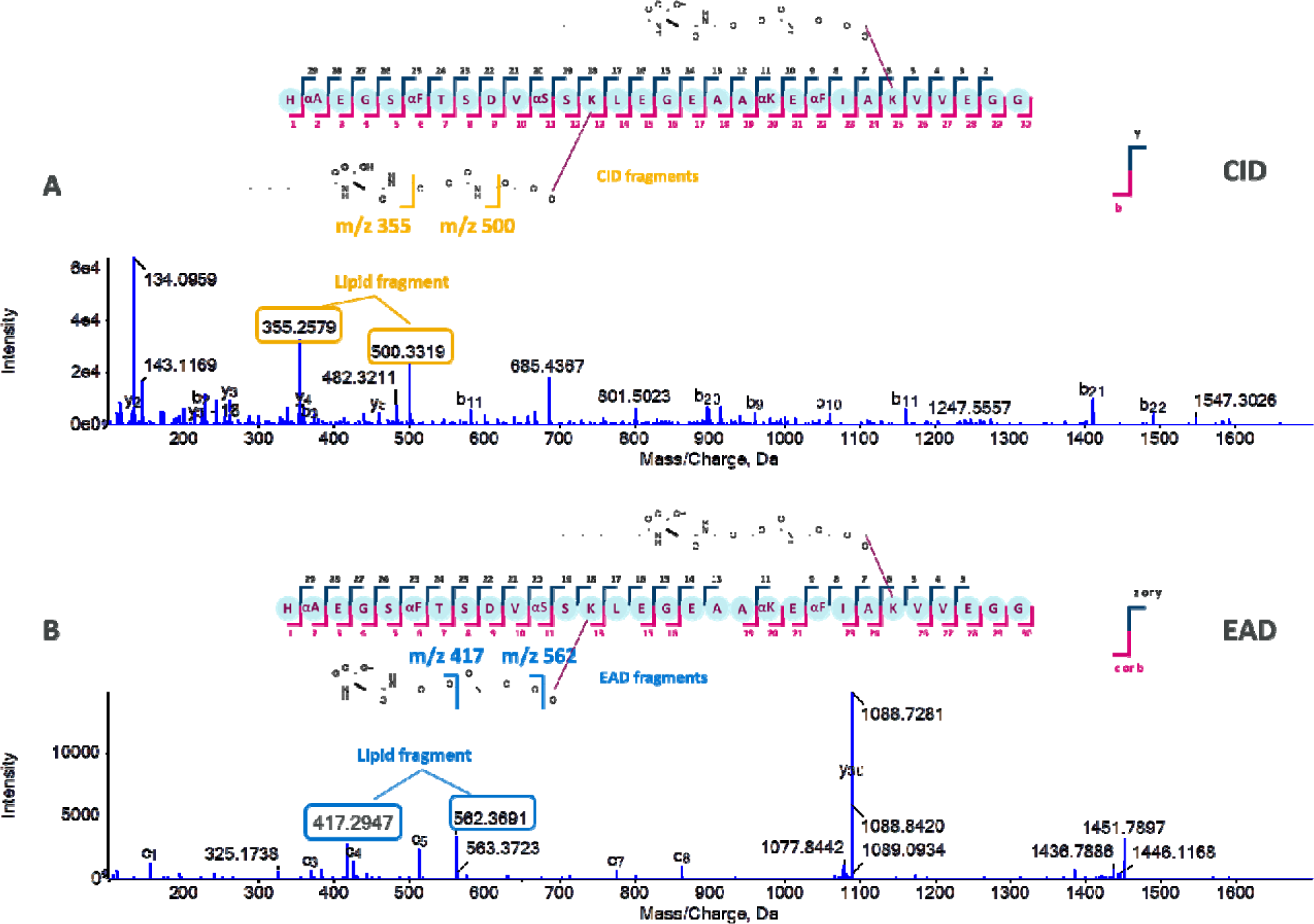
MS/MS spectra of CID (A) vs EAD (B) fragmentation of MEDI7219. “α” indicates α-L-methyl amino acid. Lipid side chain fragmentations were indicated on the structure.

**Figure 2.**
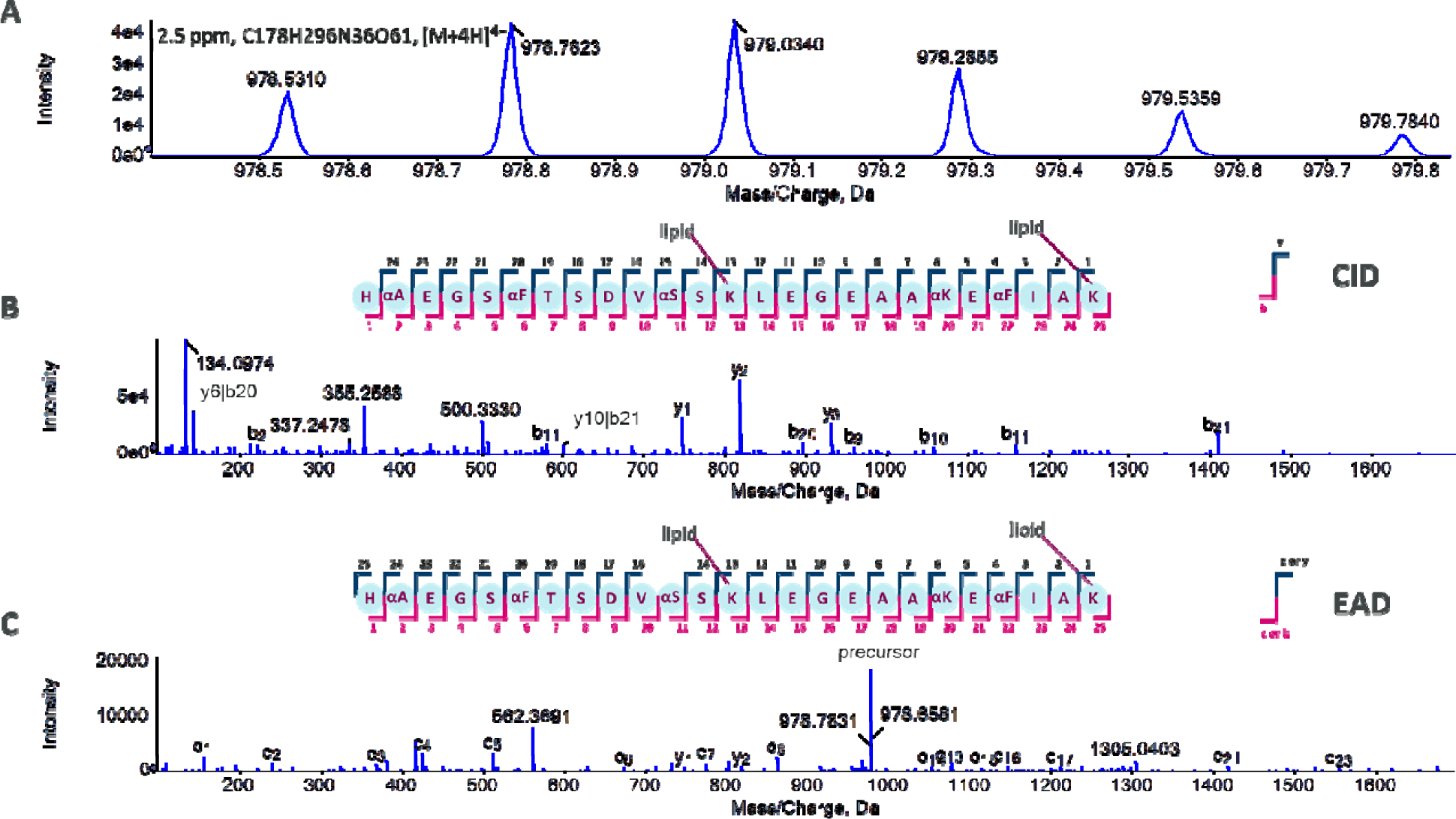
Metabolite identification example. (A) Full-scan MS showing precursor ion at m/z 978.5310. MS/MS fragmentation evidence using (B) CID and (C) EAD confirming identify of metabolite M2.

### Relative quantification of metabolites after enzymatic incubation

Metabolite chromatograms capture changes of all metabolites over pancreatin incubation period (Figure 3). Metabolite chromatograms are summed XIC (extracted ion chromatogram) of all precursors and all isotopes that belong to each metabolite. Parent MEDI7219 peptide was still present at 0.5 h although no longer the prominent species, and it decreased significantly after 2 h. Over the time course, we observe dominant peaks being left-shifted as an indication that larger peptides are further degraded to smaller peptides by proteases.

**Figure 3.**
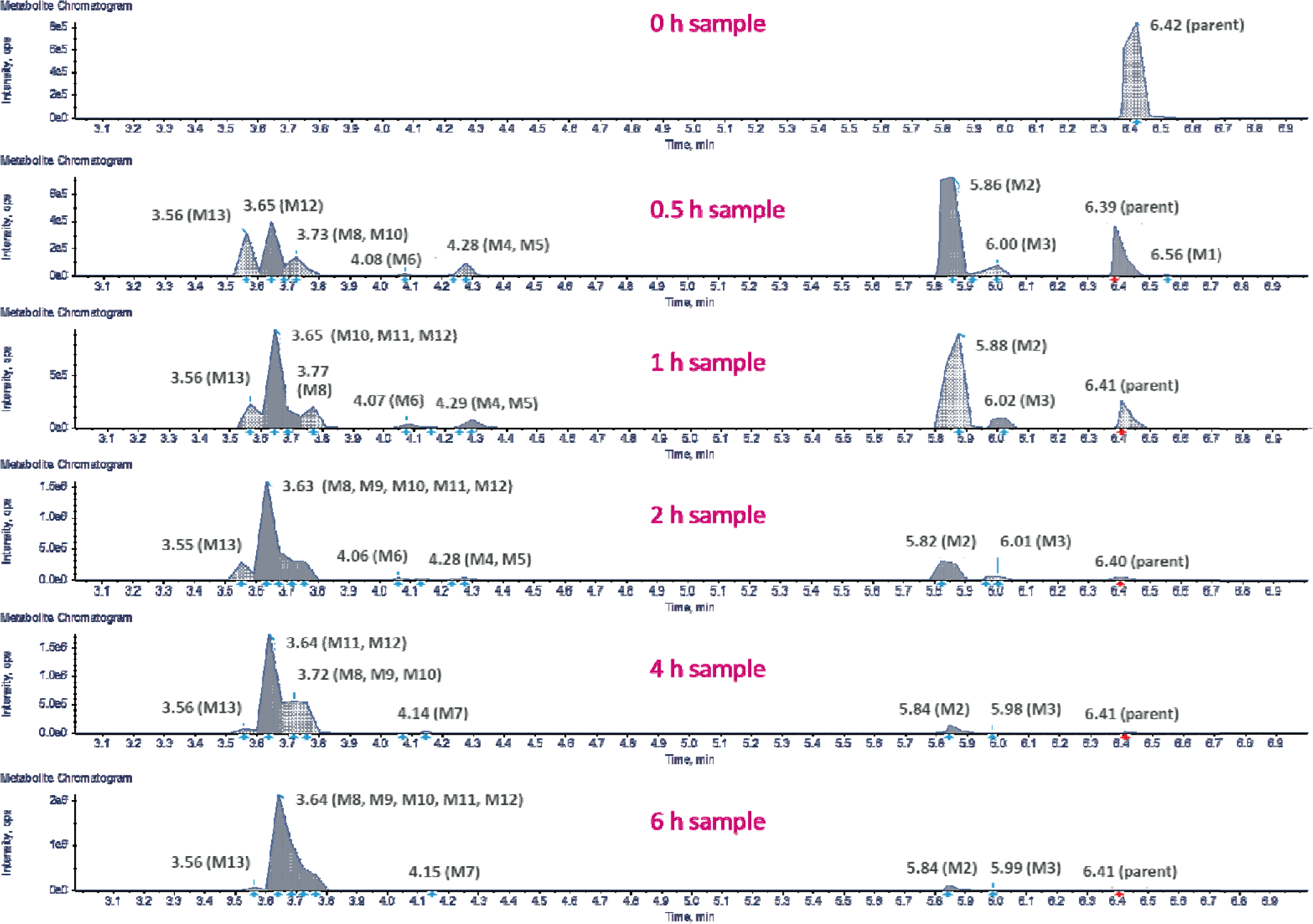
Metabolite chromatograms showing intensity changes over time (0 h, 0.5 h, 1 h, 2 h, 4 h, 6 h) from CID data

For relative quantification, after correlation analysis in MP software and exporting XIC peak areas, we calculated fractional abundance of each metabolite over the incubation period (Figure 4). This was taken as a percent of XIC peak area of each metabolite over total peak area of all metabolites at a given timepoint (Table S2). Parent peptide decreased quickly in this experiment compared to the 60 % remained intact after 2 h in previous publication (Pechenov et al., 2021). The enzyme to substrate ratio was higher in this study compared to Pechenov et al. as the assay condition was optimized to produce maximal proteolysis. It is worth noting that metabolite time profile followed 2 distinct shapes indicating 2 generations of metabolism formed. The first generation was M2 and M13 that peaked at 0.5 h and later decreased to about 2 %. The second generation of metabolites started to form later and stayed in high relative abundance over time: M12, M10 and M8. This time course trend suggests cleavage pathways where longer peptide was sequentially cleaved into smaller fragments.

**Figure 4.**
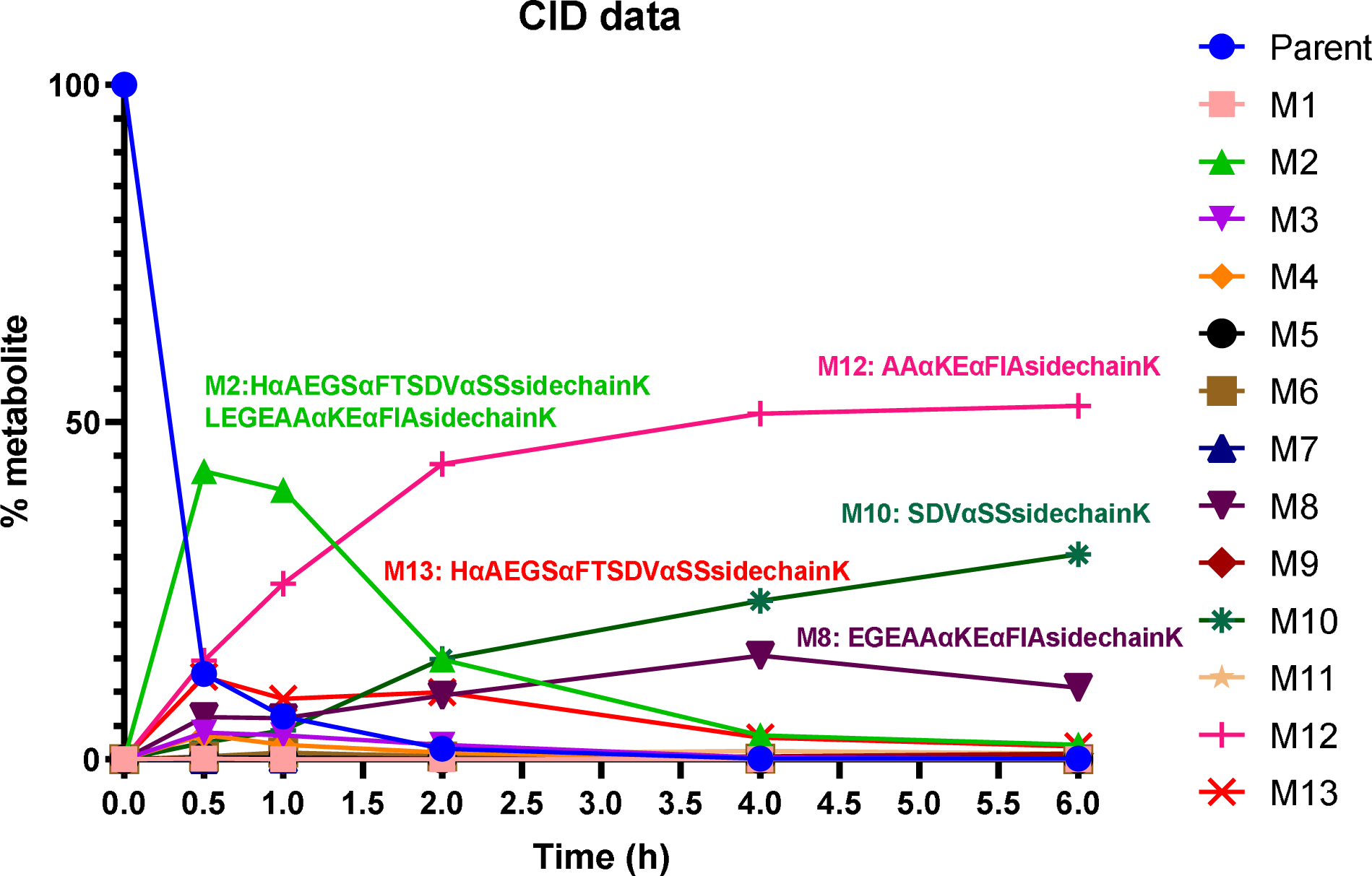
Fractional abundance calculated from individual metabolite peak area over total peak area of all peaks at a sampling timepoint. M2, M13 peaked at 0.5 h and then decreased to minor fraction. M12, M10 and M8 increased steadily overtime and stayed at high abundance.

## Discussion

In this study, both CID and EAD fragmentation methods were used to identify metabolites from a peptide with lipid side chain structure. Our results showed CID and EAD yielded same peptide identification. EAD serves as orthogonal fragmentation to CID and enhanced identification confidence.

This study identified susceptibility to pancreatic enzymes in the GLP1 agonist peptide sequence. The 13 metabolites identified map to 8 distinct cleavage sites (Figure 5). Pancreatin is a mixture of digestive enzymes including amylase, trypsin, lipase, ribonuclease and broad-spectrum protease. Trypsin cleaves C-terminal of lysine and arginine, and we identified 8 out of 13 metabolites that end with lysine. Even though the lysine is engineered with lipid side chain that may pose some steric hindrance against enzyme cleavage, this does not seem to offer complete protection. Chymotrypsin cleaves at aromatic amino acid residues such as tyrosine, tryptophan, or phenylalanine. Pepsin has similar specificity as chymotrypsin, and it preferentially cleaves at the C-terminus of phenylalanine, leucine, tyrosine and tryptophan. The 2 α-methyl-L-phenylalanines in MEDI7219 did not have cleavage products ending at those sites, so this strategy is effective against chymotrypsin and pepsin cleavage. Elastase preferentially cleaves at the C-terminus of alanine, valine, serine, glycine, leucine and isoleucine. We do observe cleavage after alanine, valine and leucine. Remaining cleavage sites observed such as C-terminal to Thr and Glu are not attributed to specific digestive protease known in pancreatin. Other enzymes in pancreatin like amylase, lipase and ribonucleases are not expected to act on this peptide structure since they break down starch, triglycerides, and RNA bond respectively.

**Figure 5.**
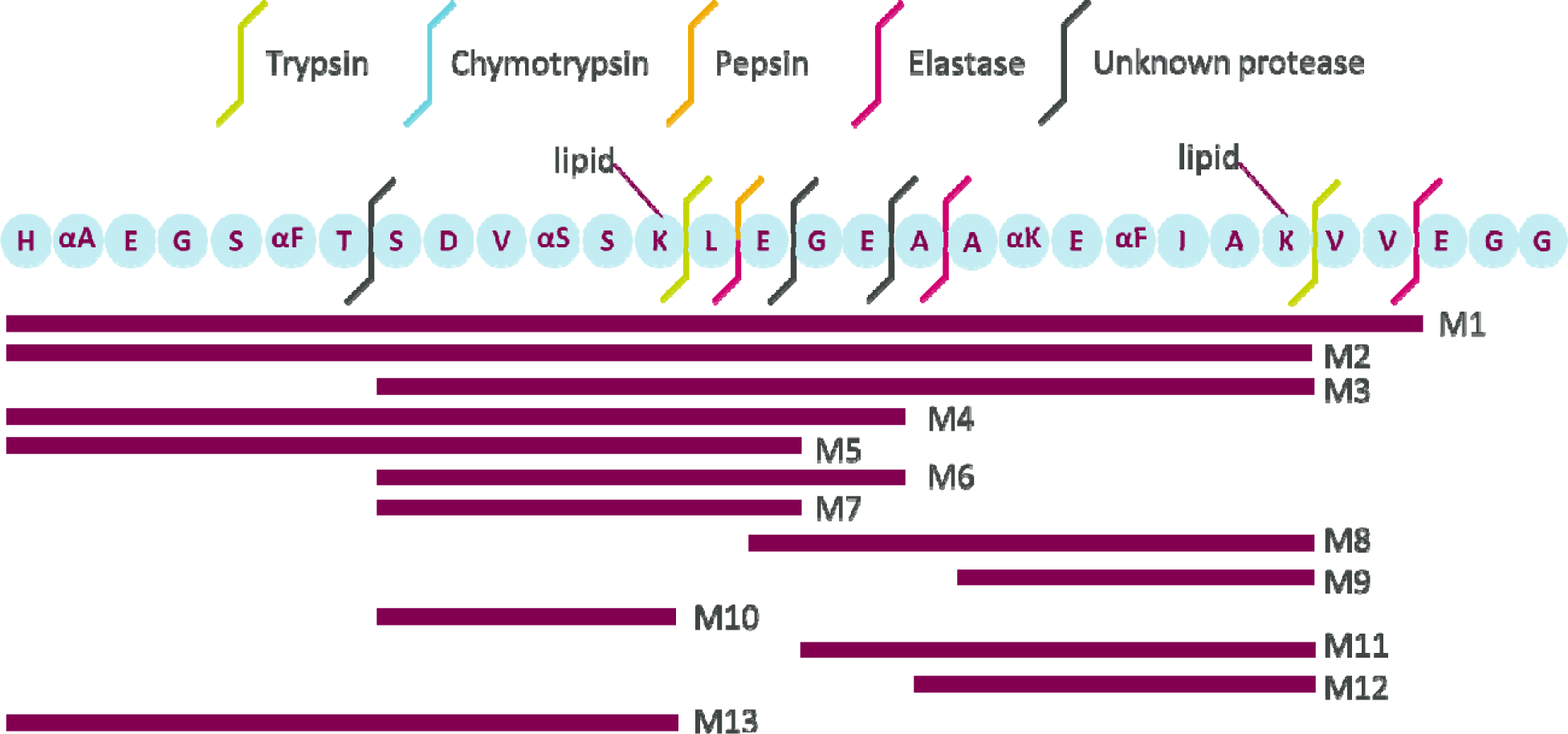
Cleavage sites identified from enzymatic incubation of MEDI7219 with pancreatin

This study expanded on previous research of proteolytic stability of oral GLP1 peptide MEDI7219 and revealed cleavage sites using pancreatin in vitro system. These proteolytically labile sites identified can guide future peptide engineering through amino acid substitution or modification to design oral drugs with better stability. Conducting in vitro soft spot analysis as demonstrated in this study offers numerous advantages over directly investigating in vivo metabolism. In vitro experiments are inexpensive and can be performed with high throughput. In addition, in vivo metabolite identification is often challenged with high matrix background from small molecules and endogenous peptides (Cuyckens et al., 2012). In vitro studies provide crucial insights for predicting metabolites observed in vivo.

## Abbreviations

CID: collision-induced dissociation
DDA: data dependent acquisition
EAD: electron activated dissociation
ECD: electron capture dissociation
ETD: electron transfer dissociation
FaSSIF: fasted-state simulated intestinal fluid
GI: gastrointestinal
GLP-1: Glucagon-like Peptide 1
KE: kinetic energy
LC-MS/MS: liquid chromatography-tandem mass spectrometry
M: metabolite
MP: Molecule Profiler
PBS: phosphate-buffered saline
XIC: extracted ion chromatogram

## Acknowledgment

We thank Haichuan Liu and Zoe Zhang from SCIEX for providing technical expertise on LC-MS instrumentation. We are also grateful for support from Yunyun Zou and Harini Kaluarachchi with SCIEX Molecule Profiler software.

## Data Availability Statement

The authors declare that all the data supporting the findings of this study are available within the paper and its Supplementary Material.

## Authorship contributions

Participated in research design: Liu, Huang, Rosenbaum.

Conducted experiments: Liu, Wang, Mu.

Performed data analysis: Liu, Wang, Mu.

Wrote or contributed to the writing of the manuscript: Liu, Huang, Wang, Mu, Rosenbaum

## Footnotes

K.L., Y.H., T.W., R.M., and A.I.R. are or were employees of AstraZeneca at the time this work was conducted and may hold stock ownership and/or stock options or interests in the company. This study was funded by AstraZeneca.

**Table 1.**
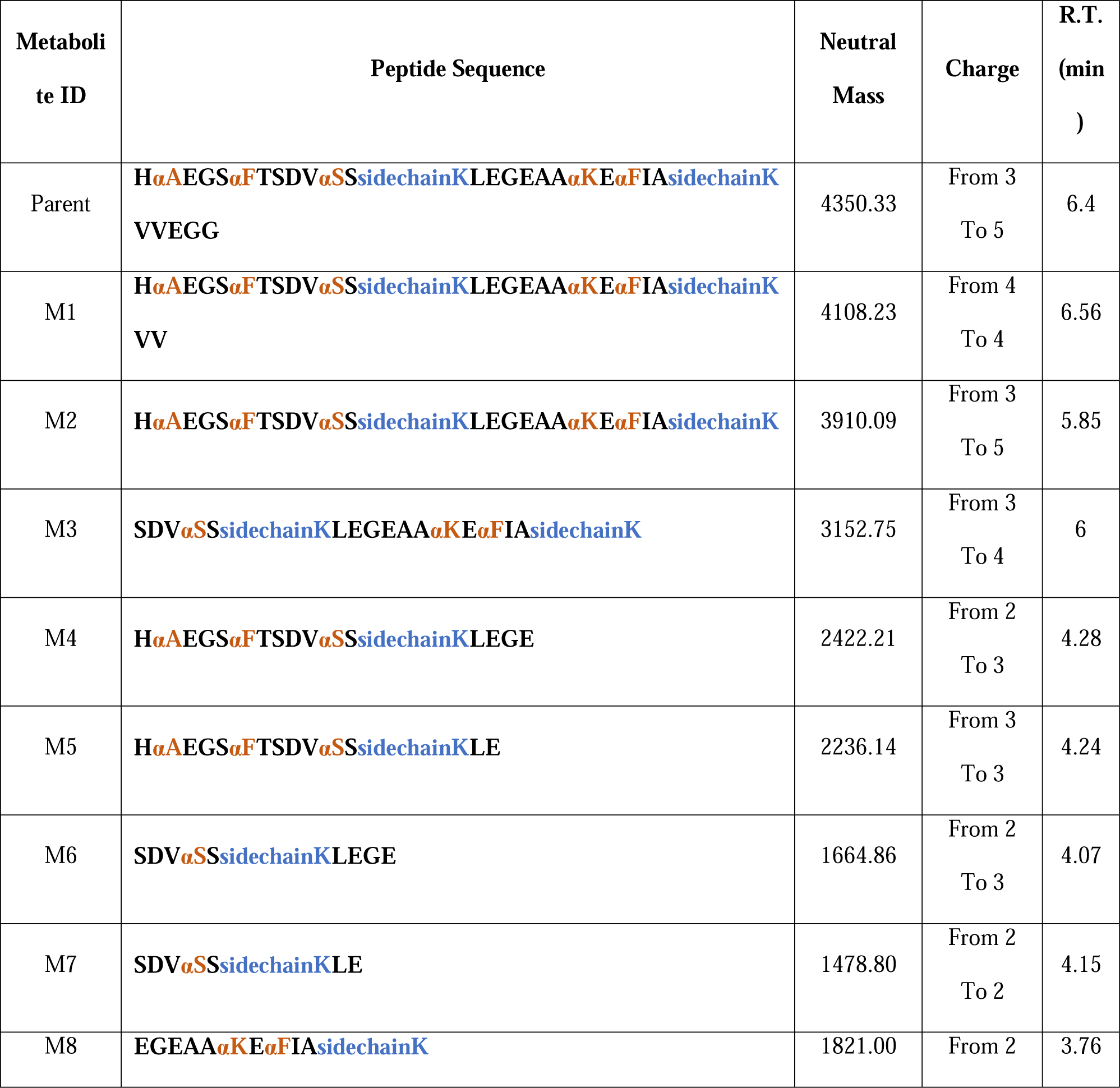
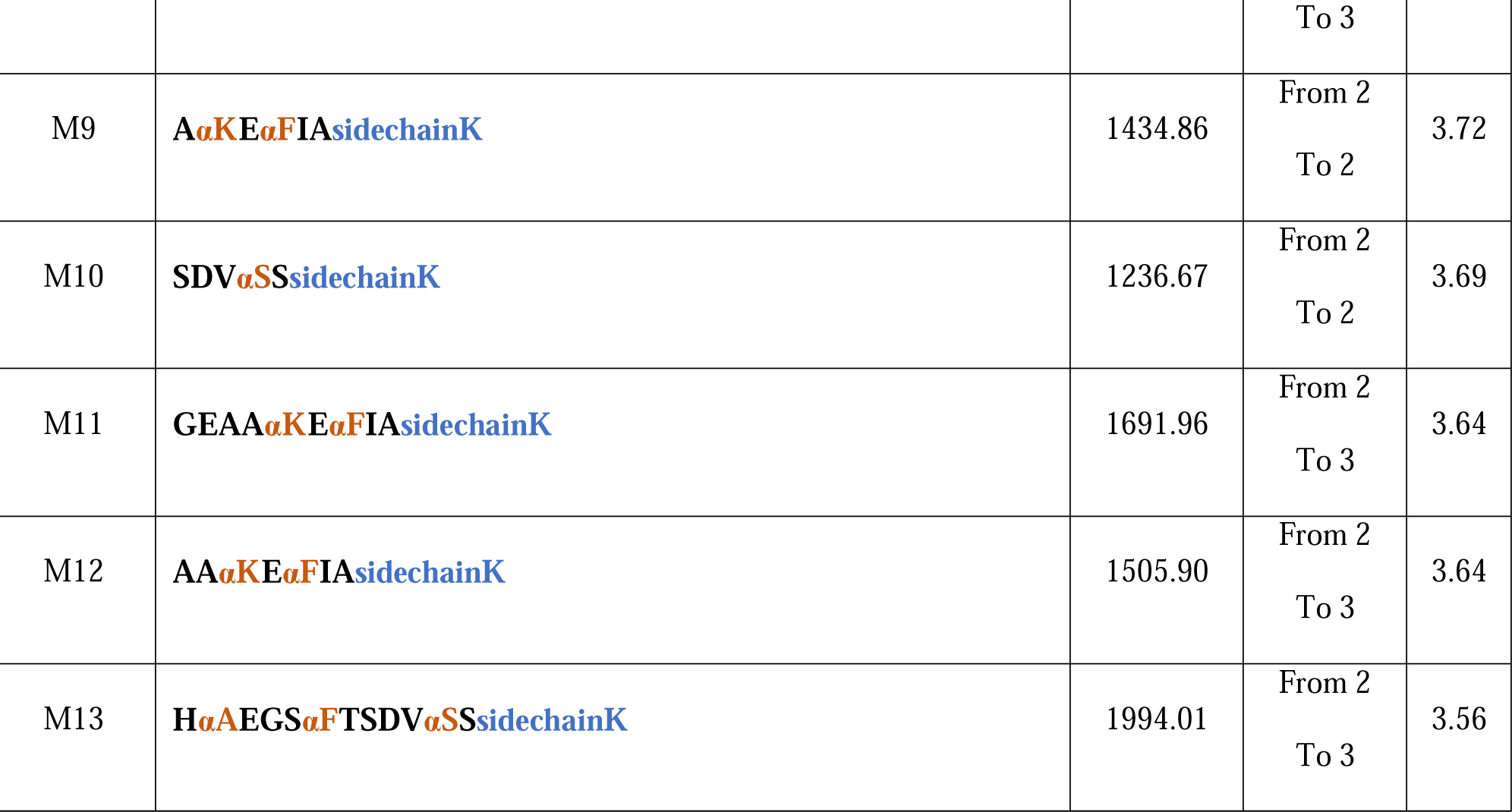
Metabolites identified from in-vitro incubation of MEDI7219 with pancreatin using either CID or EAD. In peptide sequence, "α” before an amino acid symbol indicates α-methylation (highlighted in orange) and “sidechainK” indicates lipid side chain on lysine (highlighted in blue).

## Supplementary Data

Drug Metabolism and Disposition

### EAD parameter optimization

**Figure S1.**
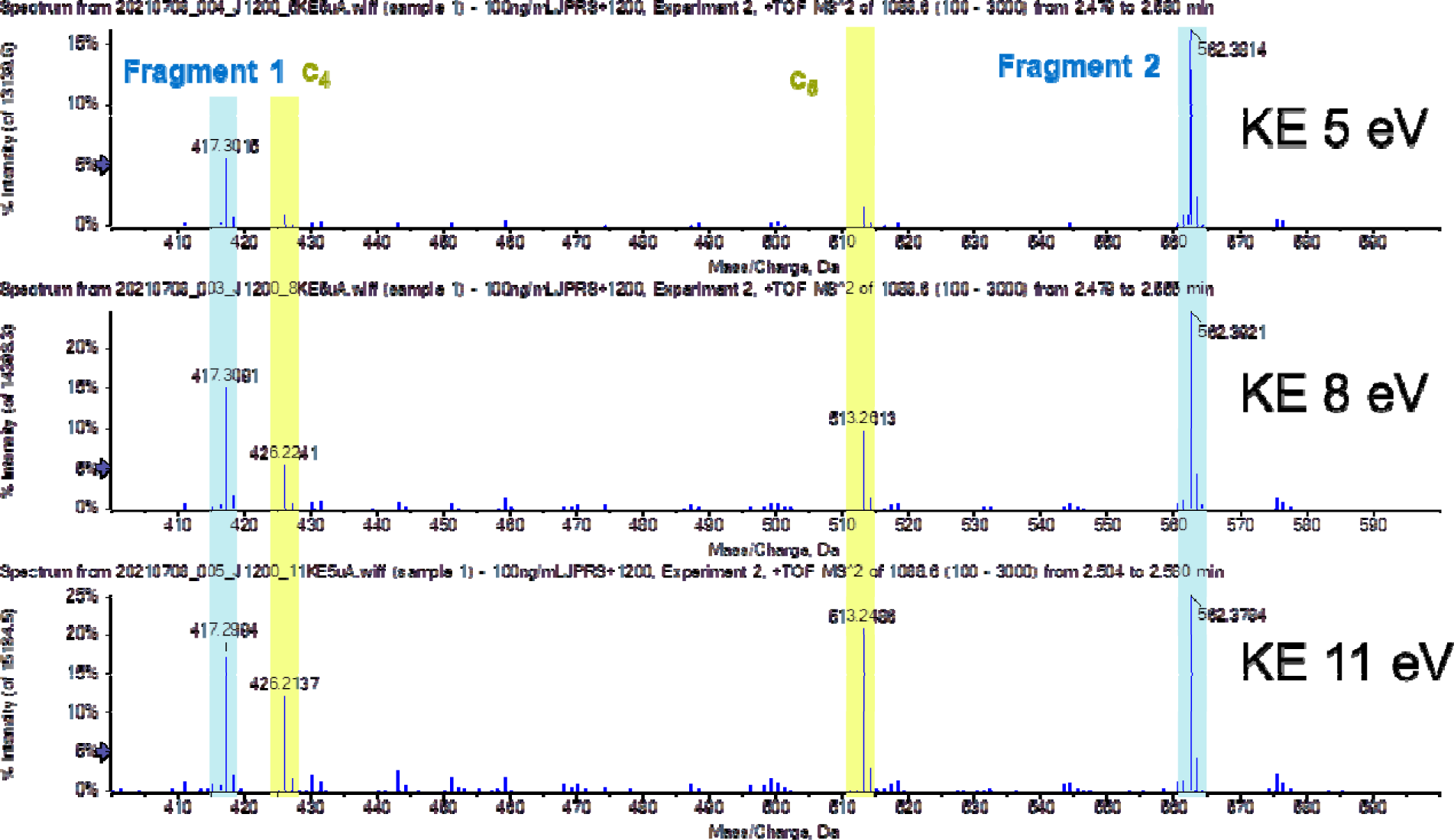
Effect of kinetic energy on EAD fragmentation. The higher the kinetic energy, the greater the relative intensity of peptide backbone fragment c4, c5 to lipid fragment 1 and 2. This indicates that cleavage of C-O on the side chain takes less energy than C-N on the backbone.

**Figure S2.**
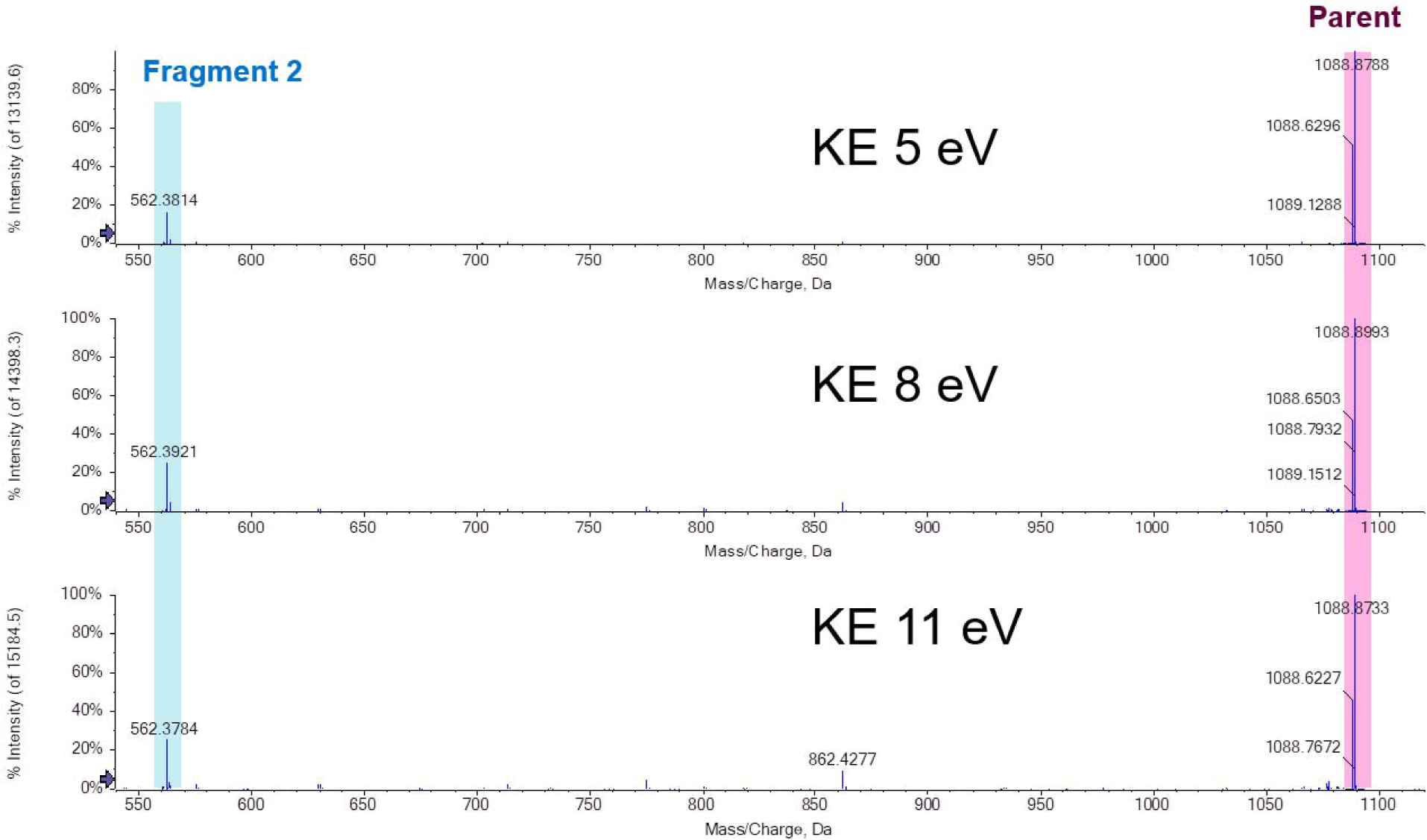
Effect of kinetic energy on EAD fragmentation. The relative intensities of lipid fragment 2 to parent ions did not change significantly with different KE

**Figure S3.**
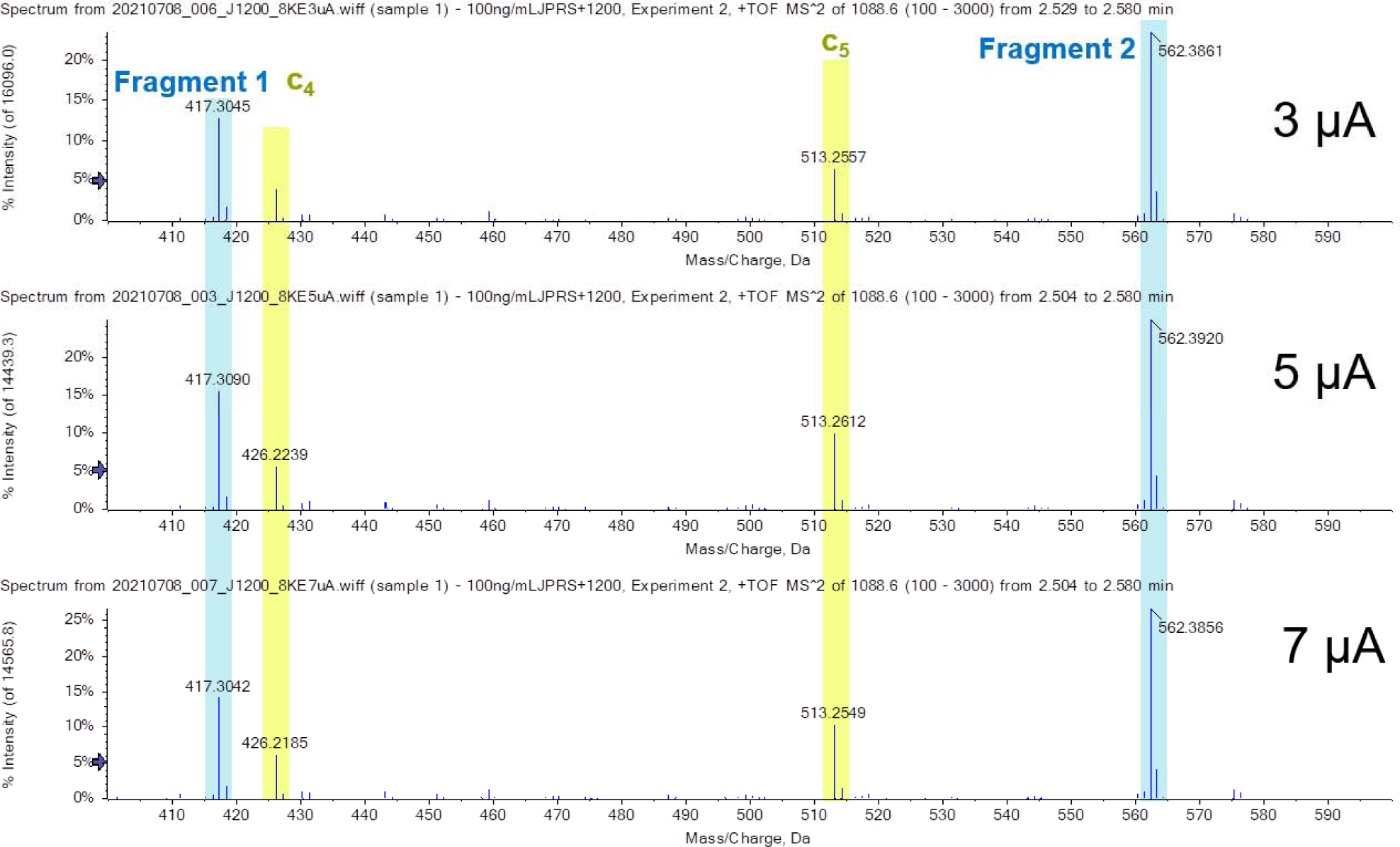
Effect of current on EAD fragmentation. The relative intensities of c4 and c5 to lipid fragment 1 and fragment 2 did not change significantly with different currents

**Figure S4.**
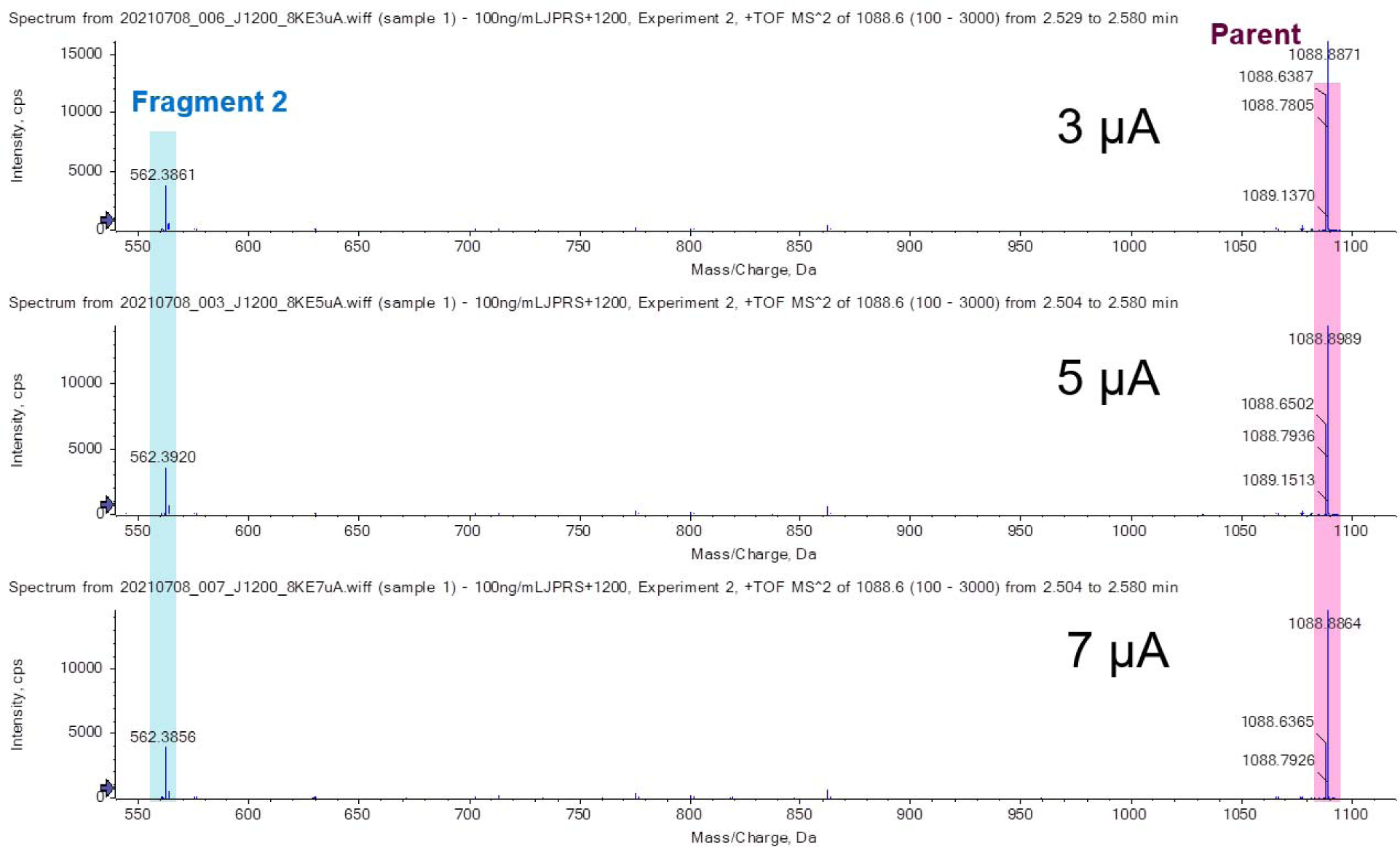
Effect of current on EAD fragmentation. The relative intensities of lipid fragment 2 to parent ions did not change significantly with different currents

**Figure S5.**
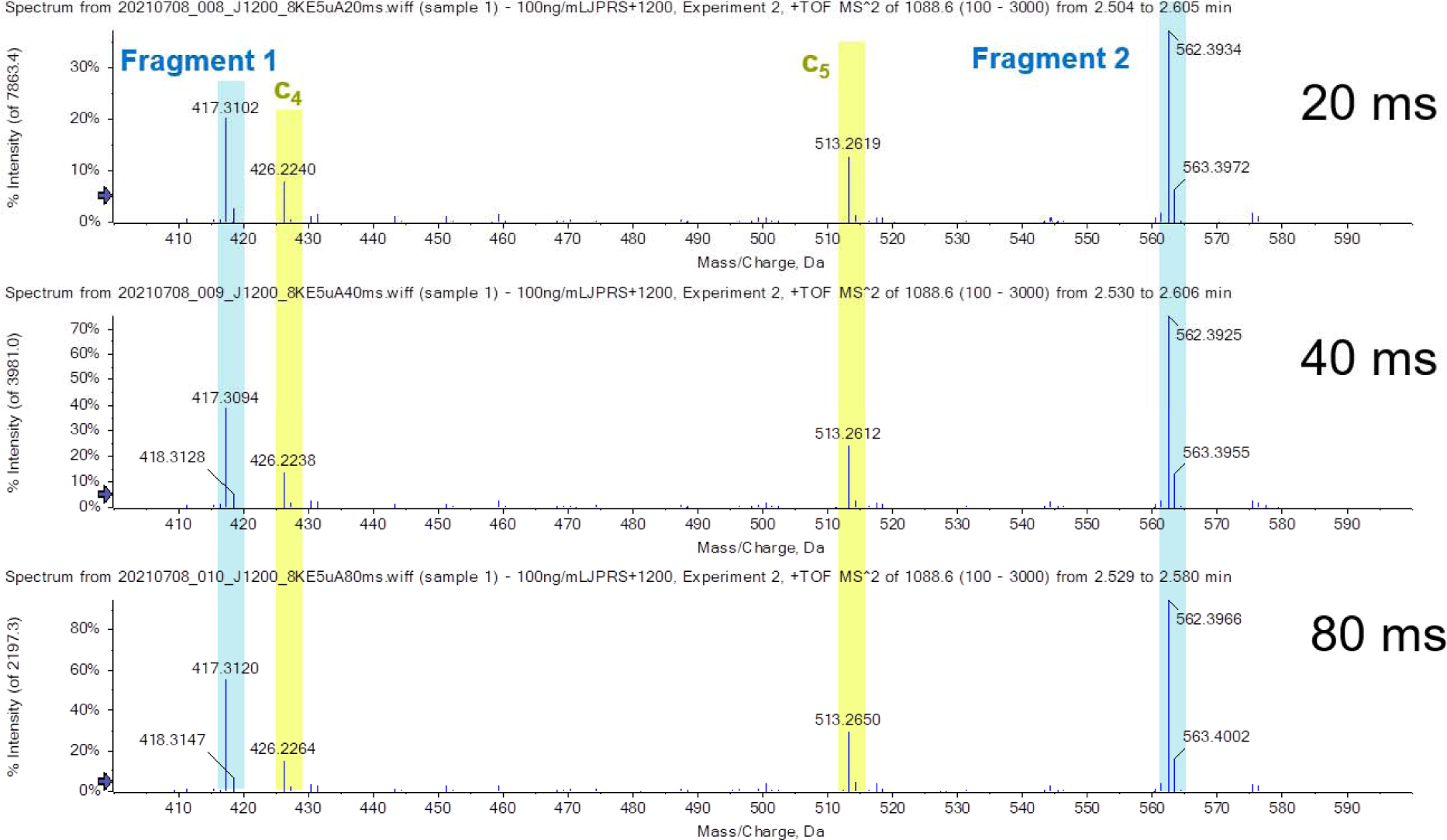
Effect of reaction time on EAD fragmentation. The relative intensities of c4 and c5 to lipid fragment 1 and fragment 2 did not change significantly with different reaction times

**Figure S6.**
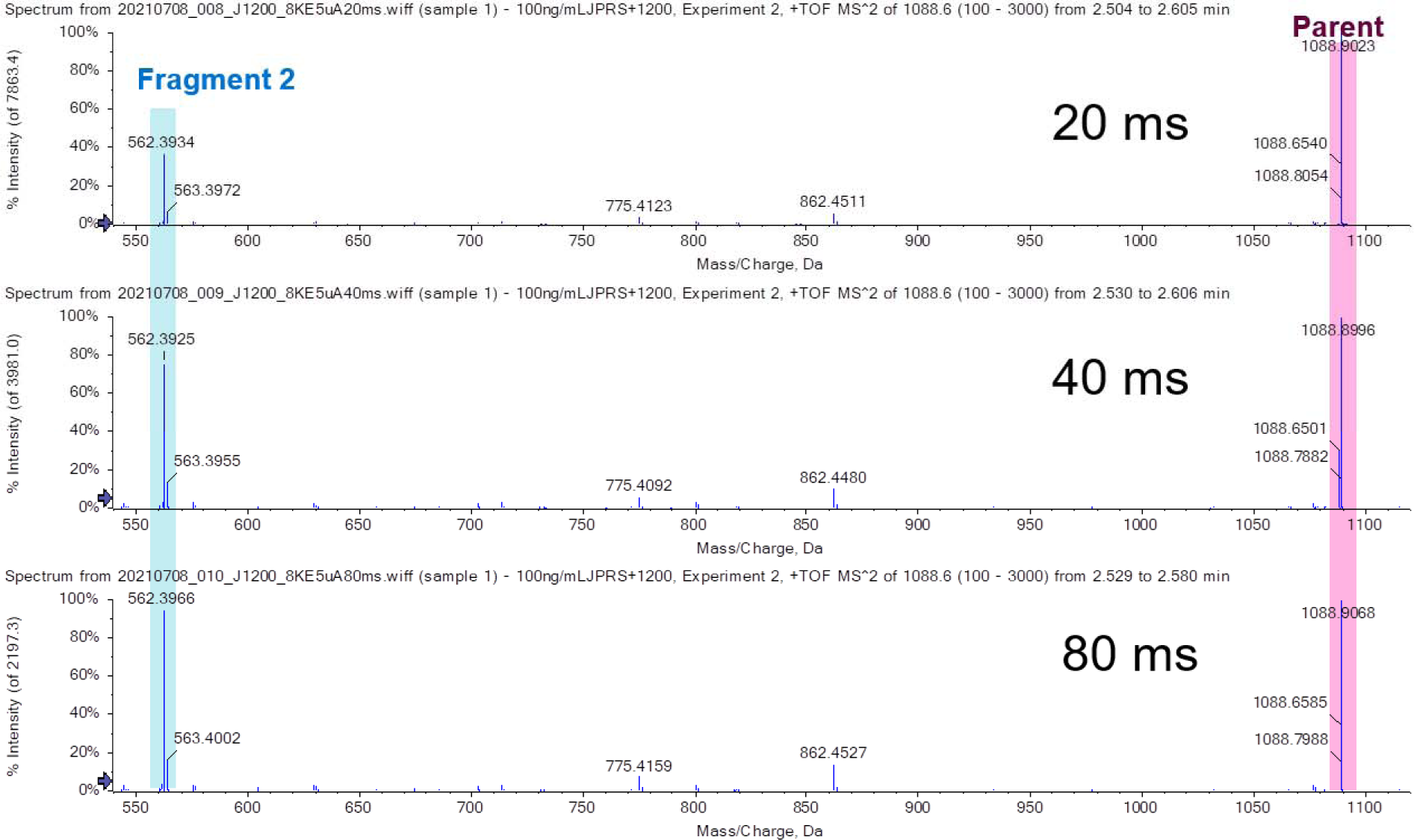
Effect of reaction time on EAD fragmentation. The longer the reaction time, the greater relative intensities of lipid fragment 2 to parent peptide. However, the absolute intensities of both fragment 2 and parent decreases as the time increases.

### Metabolite identity confirmation by LC-MS/MS

All metabolite identification evidence is shown in Figure S7-Figure S19. For each metabolite, a representative precursor with a particular charge state is chosen as an example to display MS1 spectrum and MS/MS spectrum (**Error! Reference source not found.**). The charge state was assigned based on the isotope cluster in MS1, and the structure of metabolite was confirmed based on the exact masses of precursor and product ions from CID and EAD whenever available.

**Table S1.**
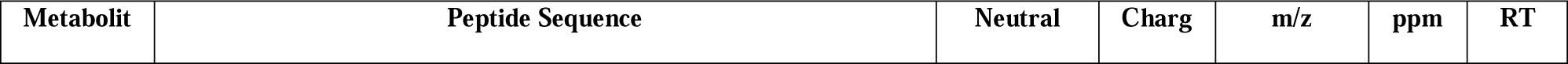
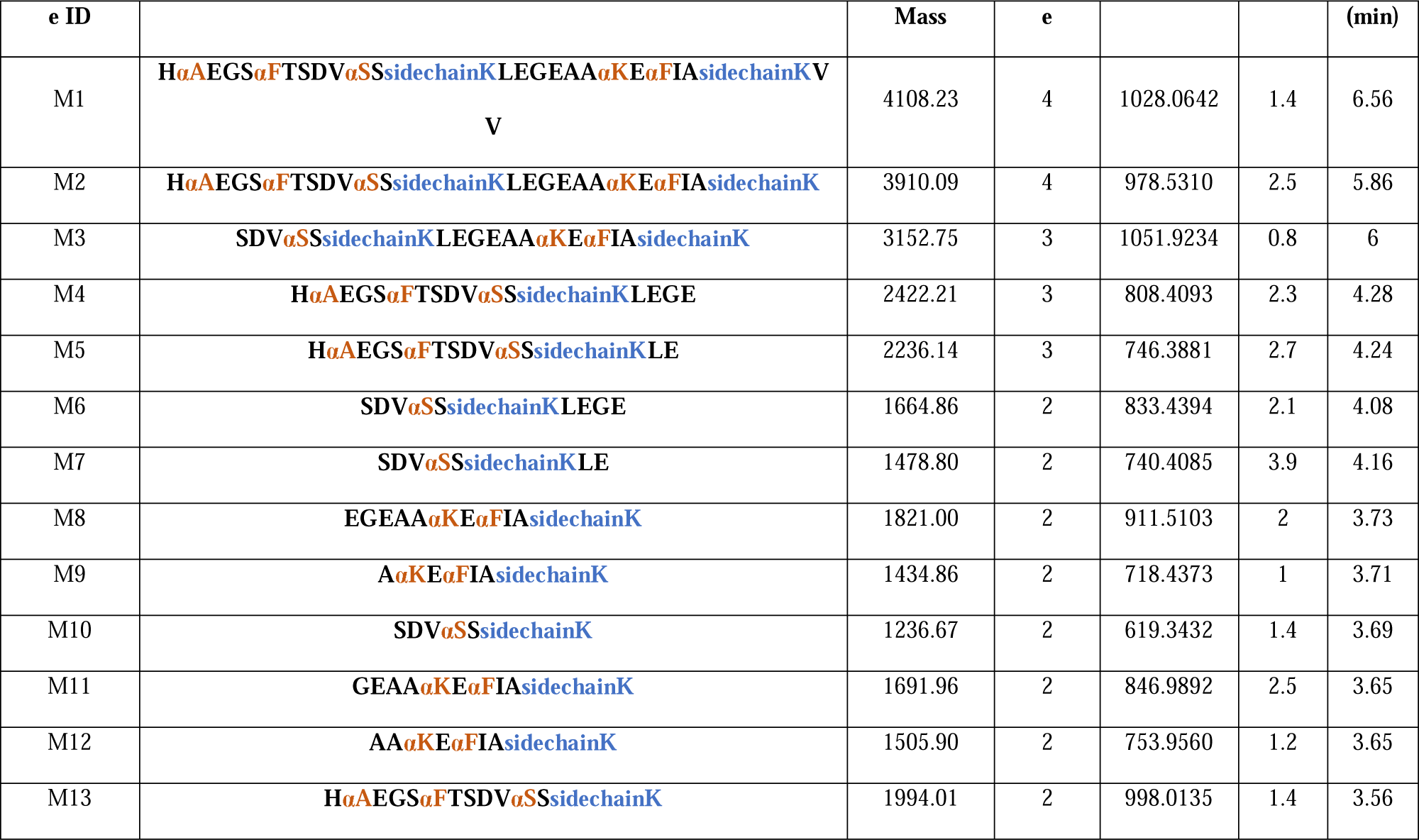
Representative precursors selected to display MS and MS/MS spectra.

**Figure S7.**
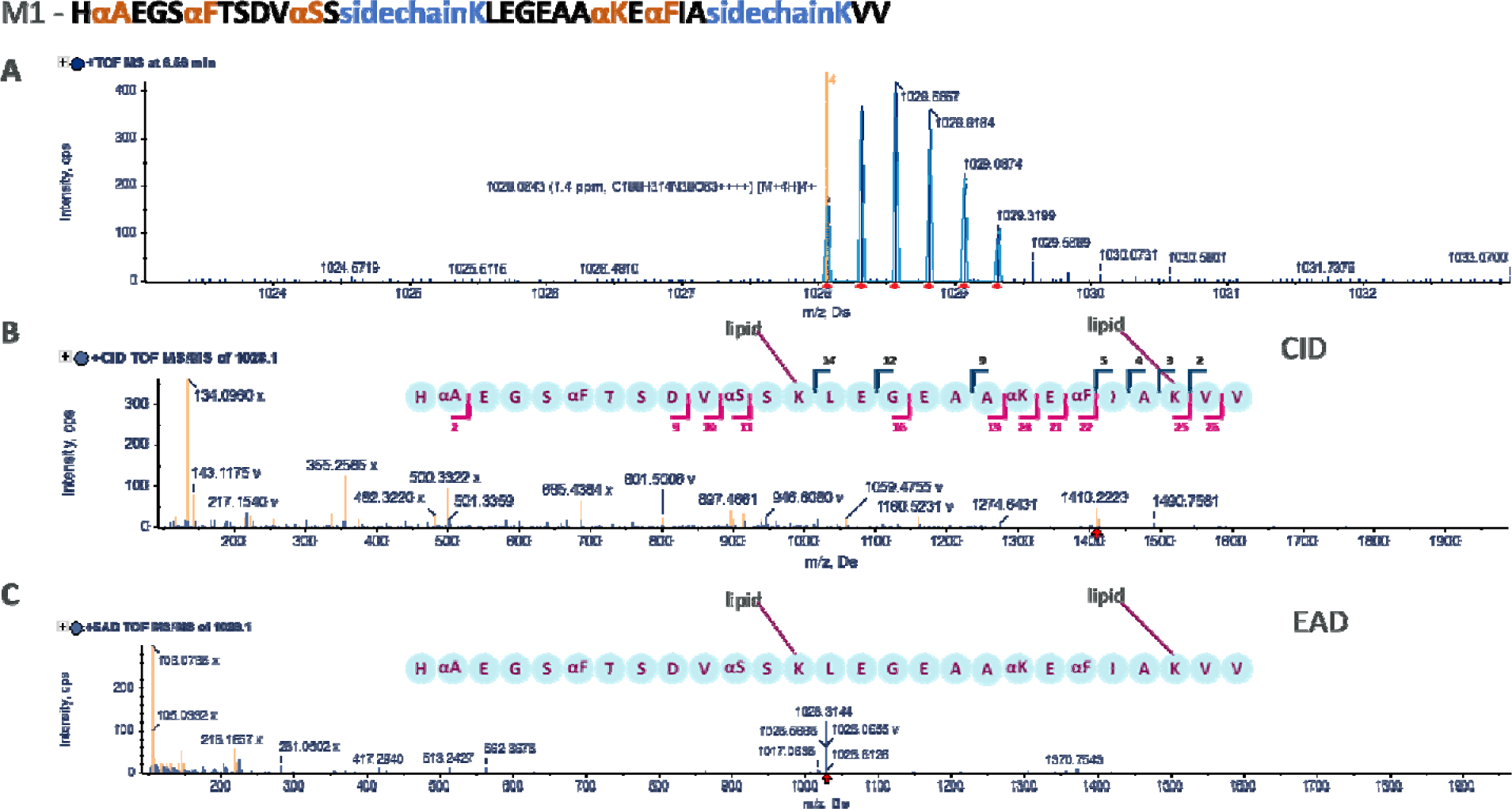
Metabolite #1 precursor isotope and fragmentation evidence

**Figure S8.**
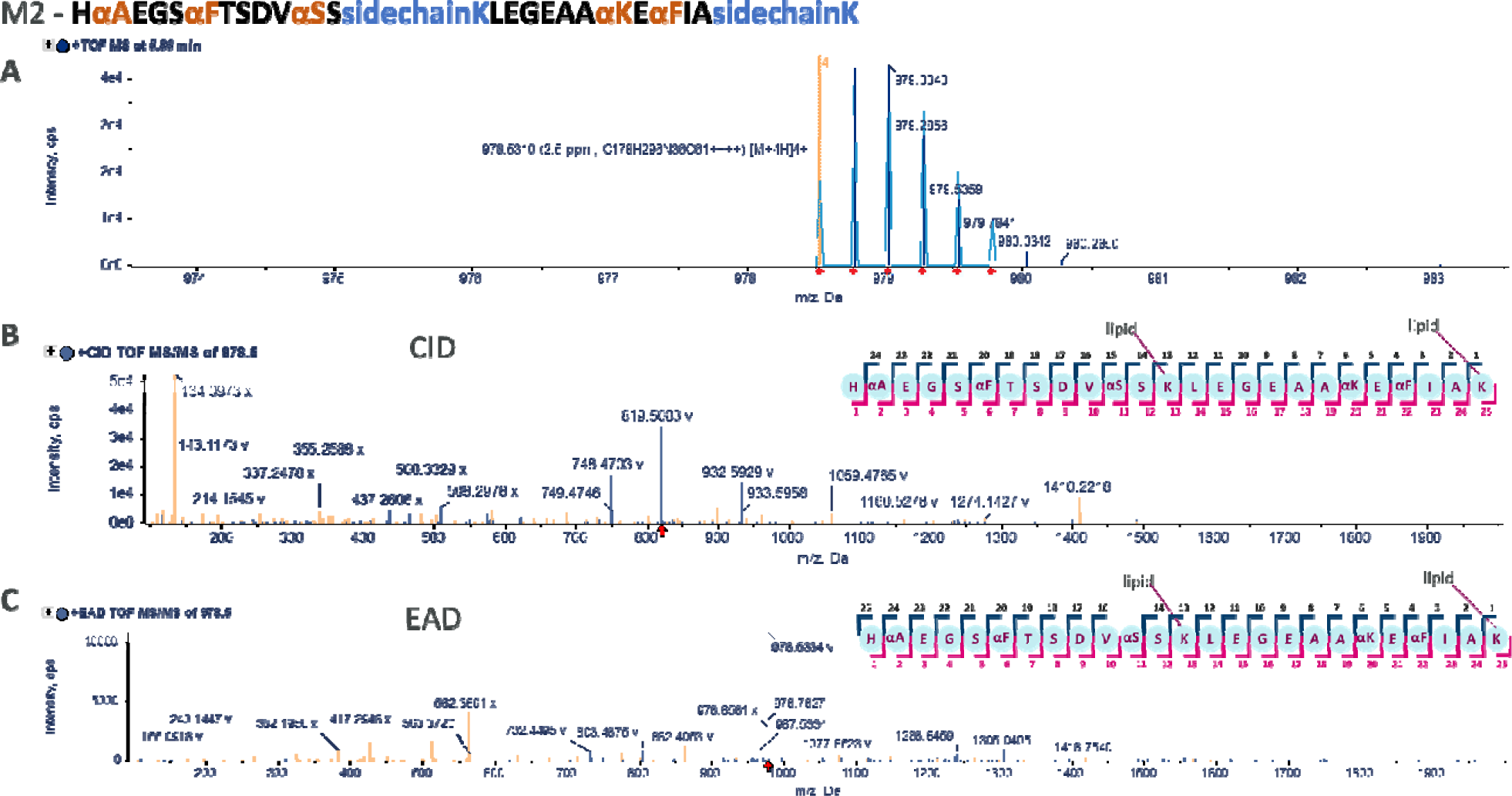
Metabolite #2 precursor isotope and fragmentation evidence

**Figure S9.**
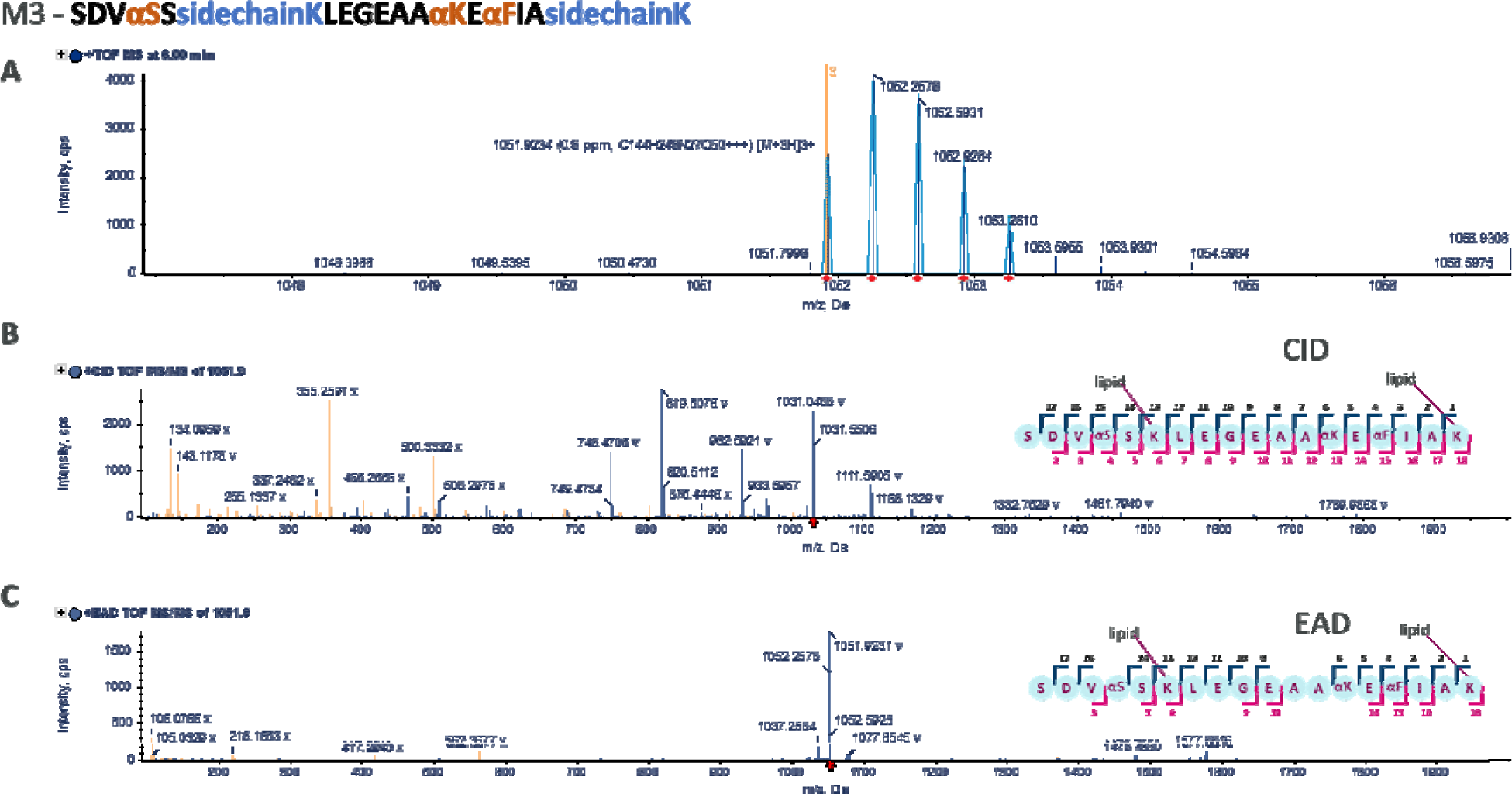
Metabolite #3 precursor isotope and fragmentation evidence

**Figure S10.**
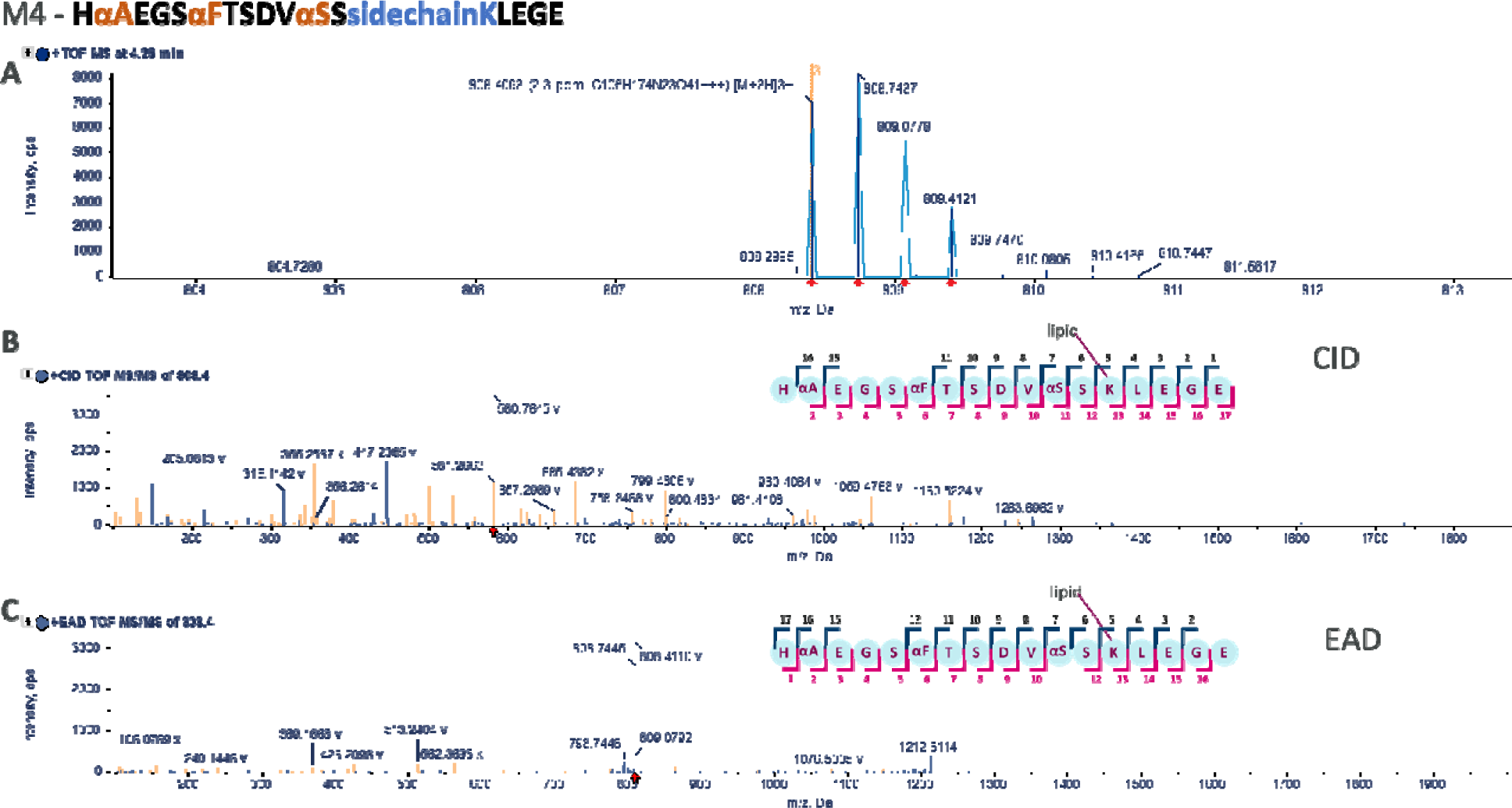
Metabolite #4 precursor isotope and fragmentation evidence

**Figure S11.**
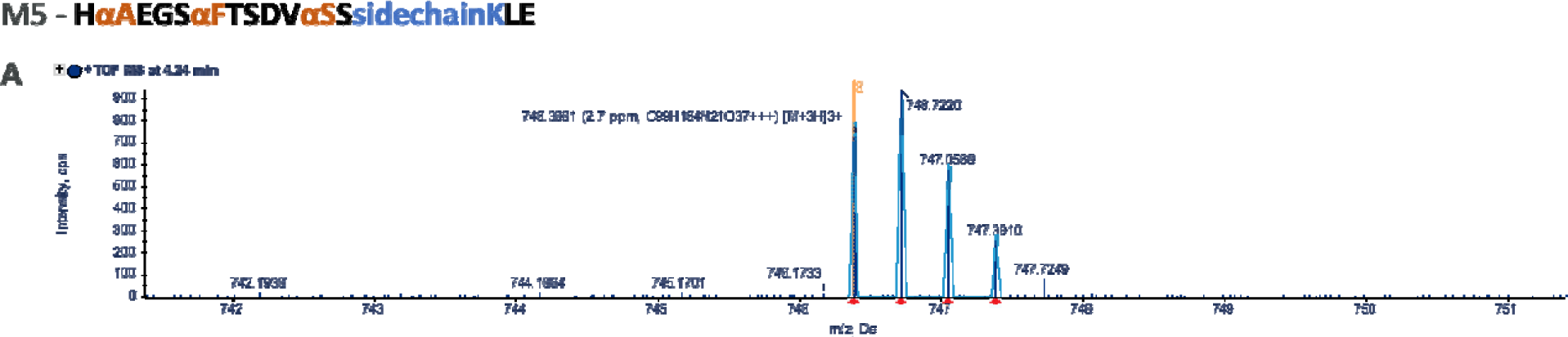
Metabolite #5 precursor isotope. No fragmentation evidence available for this metabolite

**Figure S12.**
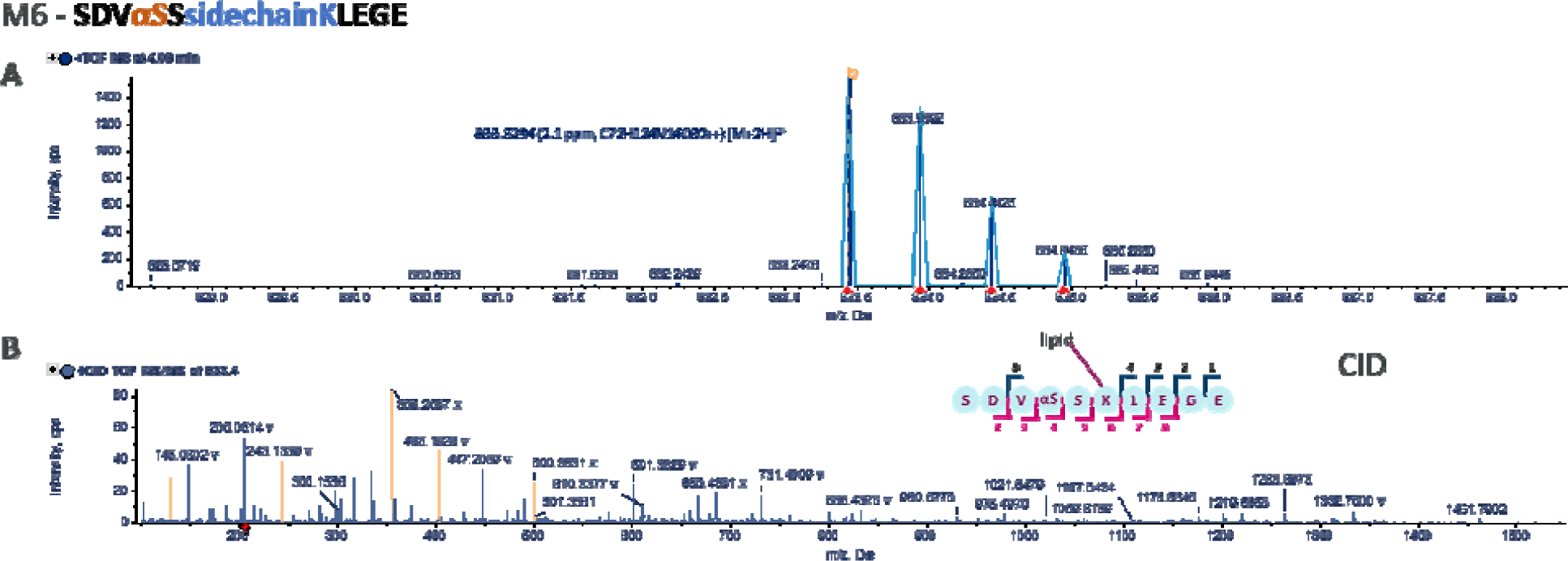
Metabolite #6 precursor isotope and fragmentation evidence. No EAD fragmentation available for this metabolite.

**Figure S13.**
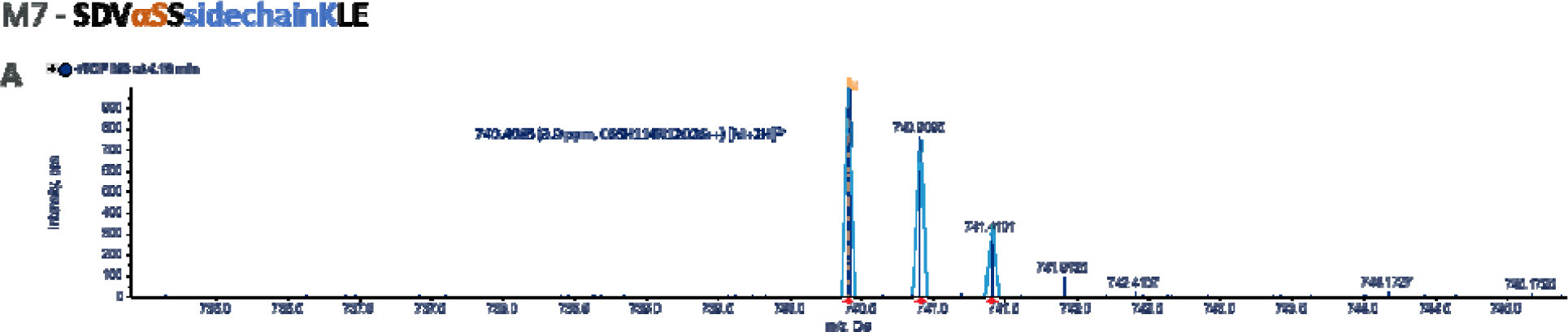
Metabolite #7 precursor isotope. No fragmentation evidence available for this metabolite

**Figure S14.**
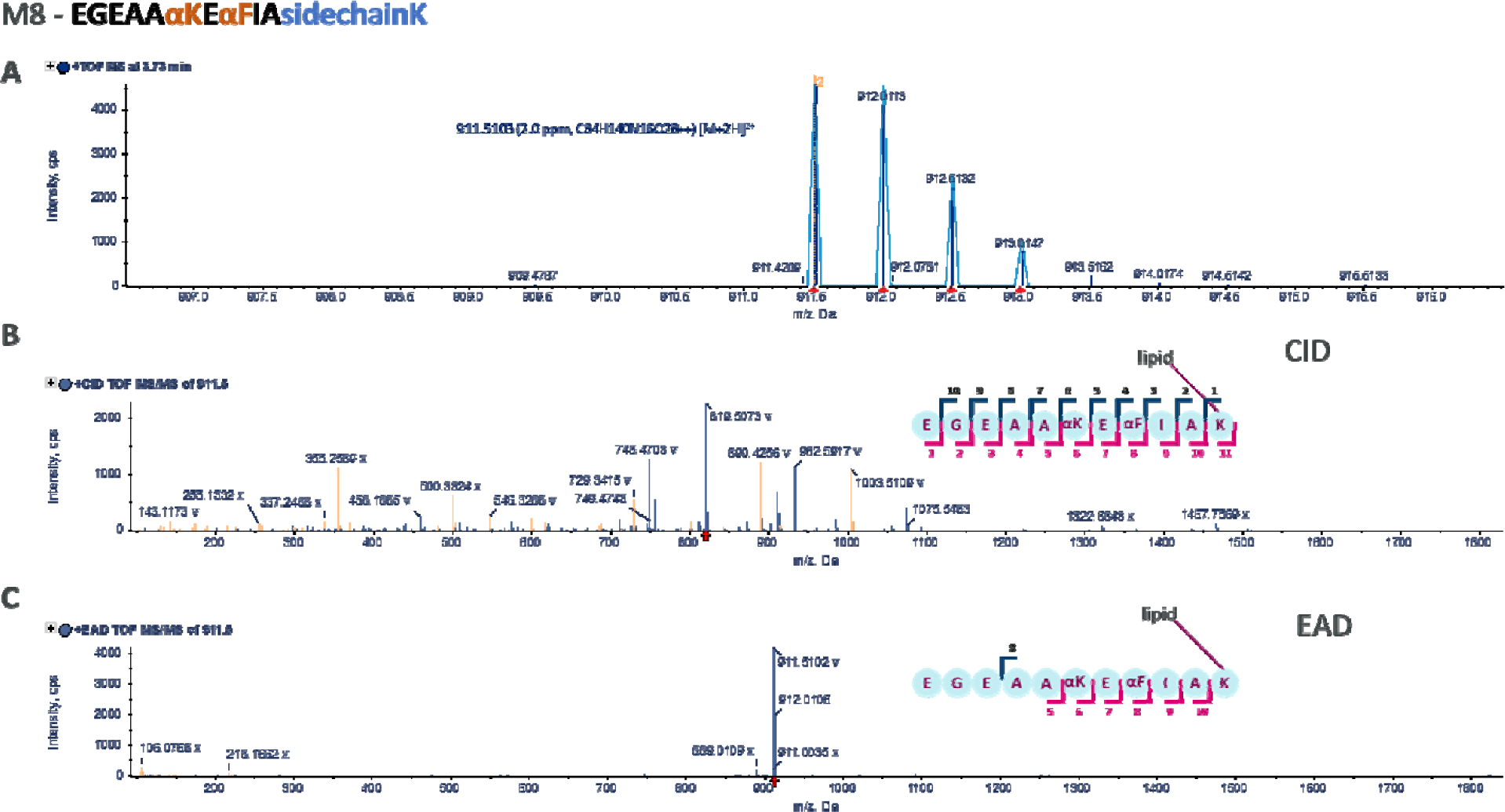
Metabolite #8 precursor isotope and fragmentation evidence

**Figure S15.**
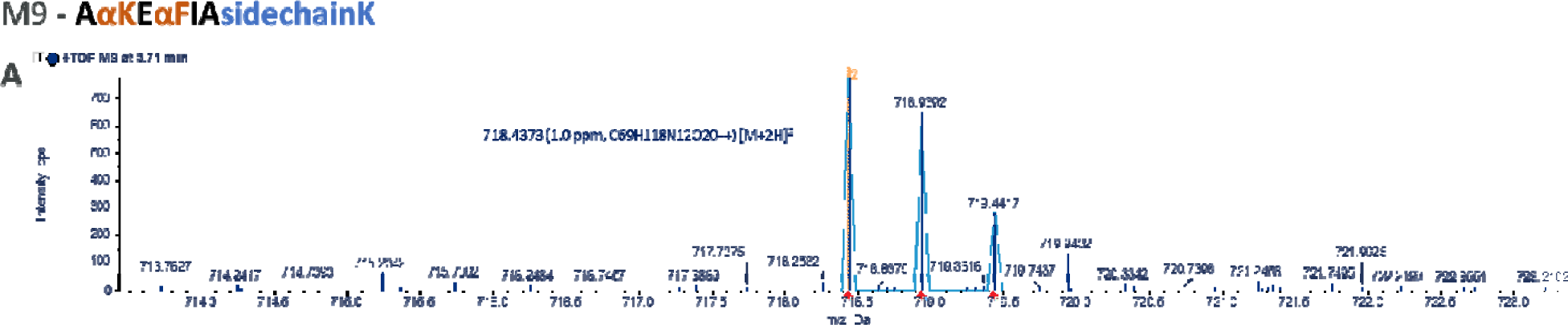
Metabolite #9 precursor isotope. No fragmentation evidence available for this metabolite

**Figure S16.**
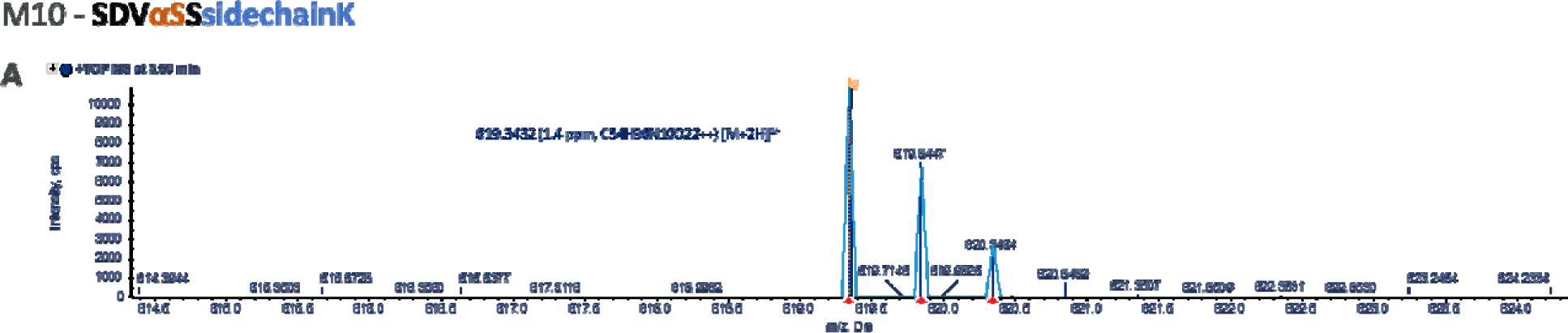
Metabolite #10 precursor isotope. No fragmentation evidence available for this metabolite

**Figure S17.**
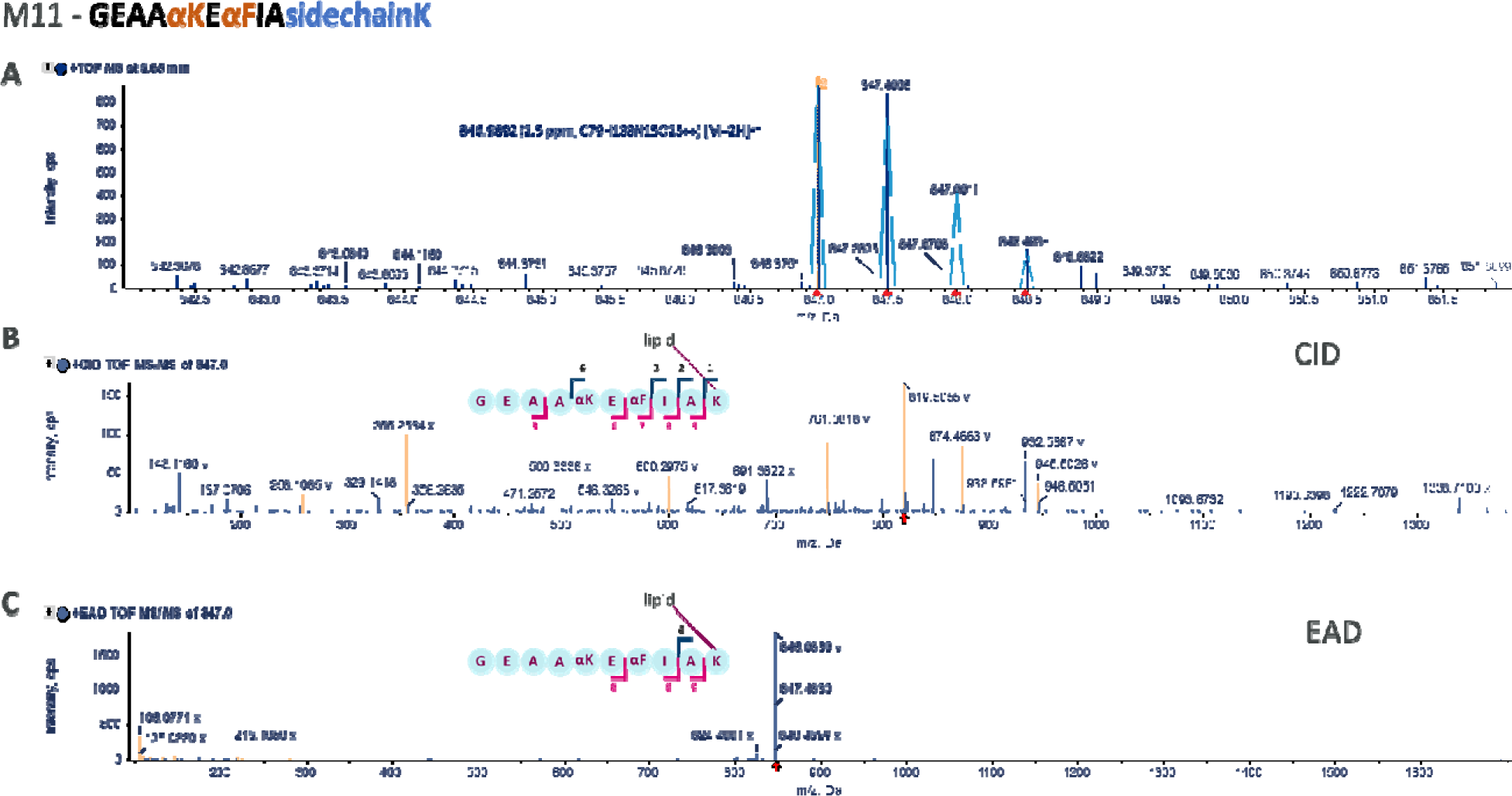
Metabolite #11 precursor isotope and fragmentation evidence

**Figure S18.**
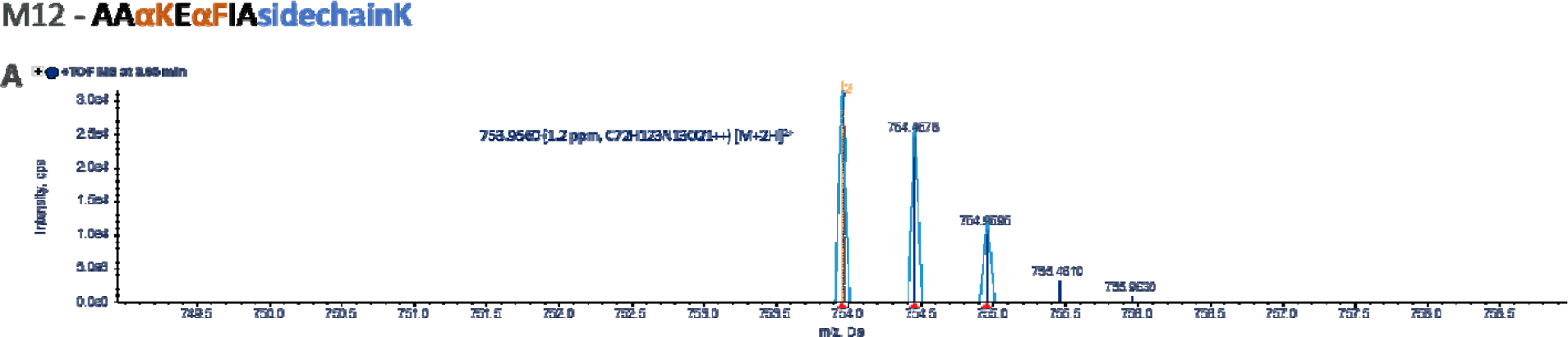
Metabolite #12 precursor isotope. No fragmentation evidence available for this metabolite.

**Figure S19.**
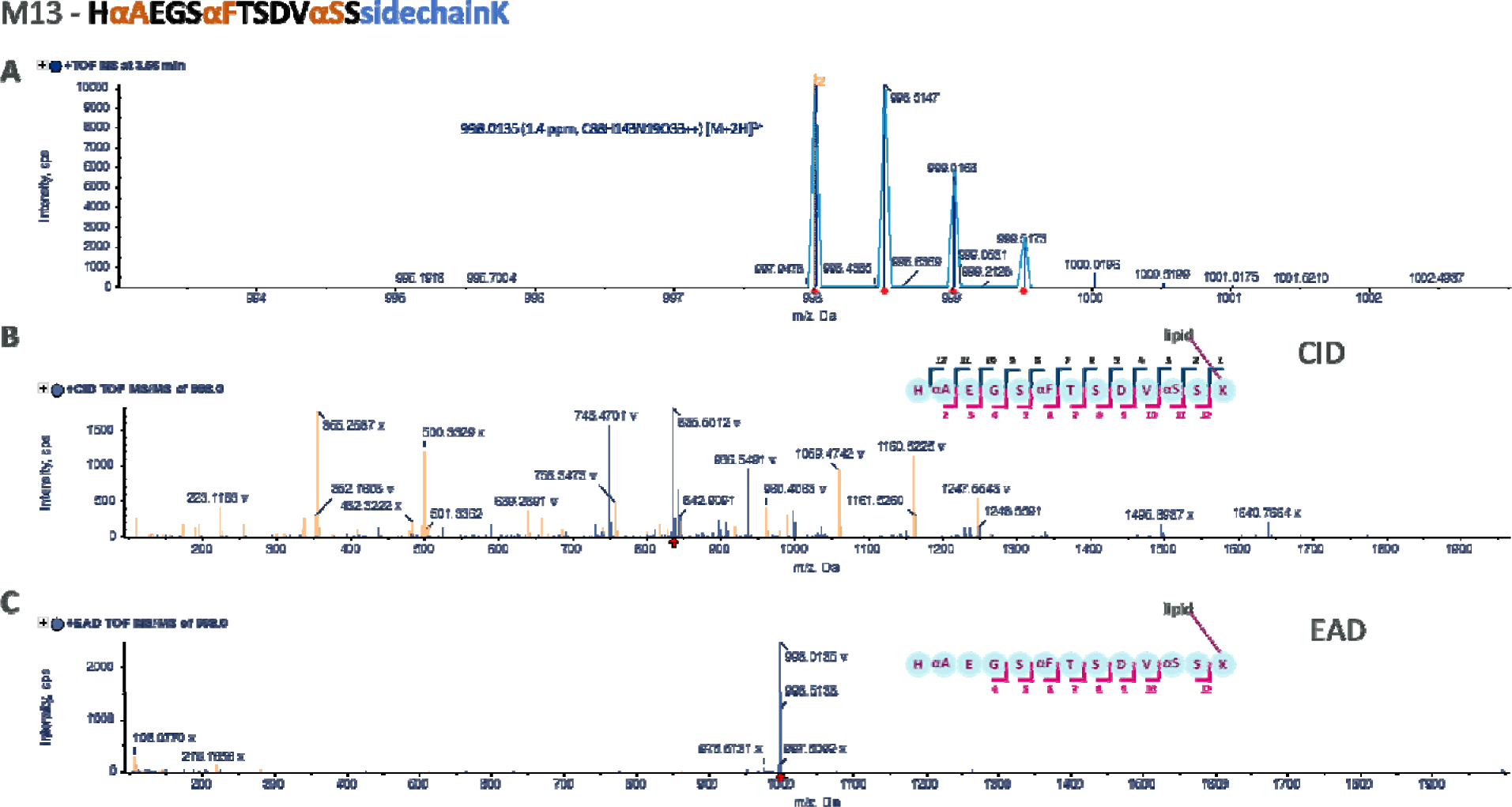
Metabolite #13 precursor isotope and fragmentation evidence

### XIC Peak Areas for All Metabolites

**Table S2.**
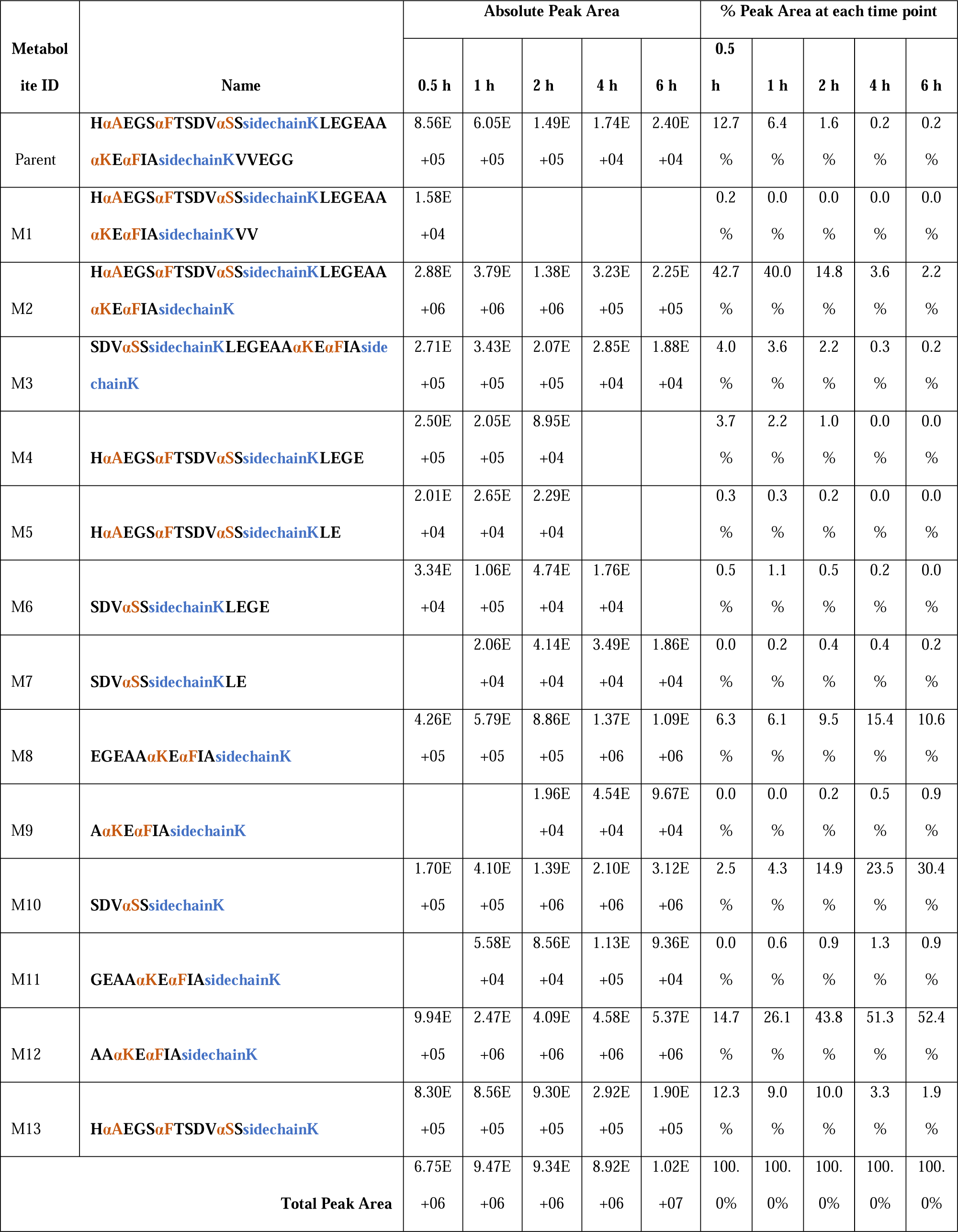
Absolute peak area of all metabolites and fractional abundance of each metabolite at each time point. The absolute peak area is a sum of all identified isotopes and charge states of each metabolite.

## References

Baba, T., Ryumin, P., Duchoslav, E., Chen, K., Chelur, A., Loyd, B., & Chernushevich, I. (2021). Dissociation of Biomolecules by an Intense Low-Energy Electron Beam in a High Sensitivity Time-of-Flight Mass Spectrometer. J Am Soc Mass Spectrom, 32(8), 1964–1975. 10.1021/jasms.0c00425

Bons, J., Hunter, C. L., Chupalov, R., Causon, J., Antonoplis, A., Rose, J., MacLean, B., & Schilling, B. (2023). Localization and Quantification of Post-Translational Modifications of Proteins Using Electron Activated Dissociation Fragmentation on a Fast-Acquisition Time-of-Flight Mass Spectrometer. J Am Soc Mass Spectrom, 34(10), 2199–2210. 10.1021/jasms.3c00144

Calabrese, V., Brunet, T. A., Degli-Esposti, D., Chaumot, A., Geffard, O., Salvador, A., Clement, Y., & Ayciriex, S. (2024). Electron-activated dissociation (EAD) for the complementary annotation of metabolites and lipids through data-dependent acquisition analysis and feature-based molecular networking, applied to the sentinel amphipod Gammarus fossarum. Anal Bioanal Chem, 416(12), 2893–2911. 10.1007/s00216-024-05232-w

Cuyckens, F., Dillen, L., Cools, W., Bockx, M., Vereyken, L., de Vries, R., & Mortishire-Smith, R. J. (2012). Identifying metabolite ions of peptide drugs in the presence of an in vivo matrix background. Bioanalysis, 4(5), 595–604. 10.4155/bio.11.333

de Haan, P., Ianovska, M. A., Mathwig, K., van Lieshout, G. A. A., Triantis, V., Bouwmeester, H., & Verpoorte, E. (2019). Digestion-on-a-chip: a continuous-flow modular microsystem recreating enzymatic digestion in the gastrointestinal tract. Lab Chip, 19(9), 1599–1609. 10.1039/c8lc01080c

Fosgerau, K., & Hoffmann, T. (2015). Peptide therapeutics: current status and future directions. Drug Discov Today, 20(1), 122–128. 10.1016/j.drudis.2014.10.003

Galia, E., Nicolaides E Fau - Hörter, D., Hörter D Fau - Löbenberg, R., Löbenberg R Fau - Reppas, C., Reppas C Fau - Dressman, J. B., & Dressman, J. B. (1998). Evaluation of various dissolution media for predicting in vivo performance of class I and II drugs. Pharm Res. (0724-8741 (Print)).

Mahapatra, M. K., Karuppasamy, M., & Sahoo, B. M. (2022). Semaglutide, a glucagon like peptide-1 receptor agonist with cardiovascular benefits for management of type 2 diabetes. Rev Endocr Metab Disord, 23(3), 521–539. 10.1007/s11154-021-09699-1

Pechenov, S., Revell, J., Will, S., Naylor, J., Tyagi, P., Patel, C., Liang, L., Tseng, L., Huang, Y., Rosenbaum, A. I., Balic, K., Konkar, A., Grimsby, J., & Subramony, J. A. (2021). Development of an orally delivered GLP-1 receptor agonist through peptide engineering and drug delivery to treat chronic disease. Sci Rep, 11(1), 22521. 10.1038/s41598-021-01750-0

Seo, K., Cho, H. W., Jeon, J. H., Kim, C. H., Lim, S., Jeong, S., Kim, K., & Chun, J. L. (2022). Influence of Bile Salts and Pancreatin on Dog Food during Static In Vitro Simulation to Mimic In Vivo Digestion. Animals (Basel), 12(20). 10.3390/ani12202734

Tyagi, P., Patel, C., Gibson, K., MacDougall, F., Pechenov, S. Y., Will, S., Revell, J., Huang, Y., Rosenbaum, A. I., Balic, K., Maharoof, U., Grimsby, J., & Subramony, J. A. (2023). Systems Biology and Peptide Engineering to Overcome Absorption Barriers for Oral Peptide Delivery: Dosage Form Optimization Case Study Preceding Clinical Translation. Pharmaceutics, 15(10). 10.3390/pharmaceutics15102436

Tyagi, P., Trivedi, R., Pechenov, S., Patel, C., Revell, J., Wills, S., Huang, Y., Rosenbaum, A. I., & Subramony, J. A. (2021). Targeted oral peptide delivery using multi-unit particulates: Drug and permeation enhancer layering approach. J Control Release, 338, 784–791. 10.1016/j.jconrel.2021.09.002

Yang, J., Yuan, J., Huang, Y., & Rosenbaum, A. I. (2024). Reference-Free Thio-Succinimide Isomerization Characterization by Electron-Activated Dissociation. ChemRxiv. 10.26434/chemrxiv-2024-9p6j7

Yu, X., Fridman, A., Bagchi, A., Xu, S., Kwasnjuk, K. A., Lu, P., & Cancilla, M. T. (2020). Metabolite Identification of Therapeutic Peptides and Proteins by Top-down Differential Mass Spectrometry and Metabolite Database Matching. Anal Chem, 92(12), 8298–8305. 10.1021/acs.analchem.0c00652

